# Sequence-encoded interactions program internal condensate architecture

**DOI:** 10.1101/2025.10.02.680069

**Authors:** Sumit Majumder, Ashif Akram, Angelina Ning, Jeremy Schmit, Ankur Jain

## Abstract

Many cellular condensates, such as the nucleolus and stress granules, contain multiple coexisting phases with distinct compositions and material properties. This internal organization is crucial for function, yet how it is established remains unclear. Here, using a programmable DNA system, we reveal how molecular interactions can precisely encode multiphase architecture. We find that phase separation drives macromolecules into a semi-dilute regime where subtle differences in homotypic interaction energies are amplified into dominant organizational forces. A critical interaction energy threshold must be overcome to trigger internal demixing, after which molecular partitioning scales near-linearly with interaction strength. This universal relationship is captured by an associative polymer model, and enables engineering of condensates with up to four coexisting phases exhibiting 100-fold differences in viscosity within the same droplet. These design principles extend to RNA-peptide systems, establishing a general framework for how sequence can program hierarchical self-assembly and organize biological matter.

## Introduction

Eukaryotic cells exhibit a remarkable degree of internal organization, much of which is achieved without the use of traditional membrane-bound organelles. A key principle underlying this compartmentalization is the formation of biomolecular condensates^1–3^. These condensates are dynamic assemblies of proteins and nucleic acids that arise via phase separation from the surrounding nucleoplasm or cytoplasm^4–6^. By concentrating specific biomolecules, condensates create microenvironments that orchestrate complex cellular processes and, when dysregulated, contribute to pathological states such as neurodegeneration and cancer^7–12^.

Over the past decade, significant strides have been made in understanding the principles governing condensate formation. Studies across proteins, nucleic acids, and synthetic polymers have revealed how sequence-encoded features, such as interaction valency^5,12–14^, charge patterning^15,16^, and structural disorder^17,18^, provide the driving force for macromolecules to phase separate into liquid-like states. These discoveries are allowing us to put together a framework that links molecular sequence to phase behavior, enabling predictions of when condensation occurs, which molecules will be enriched, and how the properties of the resulting condensate can be tuned^19–21^. This molecular grammar also explains how condensates create selective biochemical environments, transforming our understanding of cellular biochemical organization and suggesting avenues for therapeutic intervention^10,22^.

However, many cellular condensates are far more complex than simple homogeneous droplets and exhibit sophisticated internal architectures with multiple coexisting phases. The nucleolus, for example, contains at least three distinct sub-compartments: the fibrillar center, dense fibrillar component, and granular component, each enriched with specific proteins that faithfully localize to their designated regions^6,23,24^. Similarly, stress granules display core-shell architectures where stable core proteins remain segregated from the dynamic shell components^8^. On the other hand, several condensates appear partially miscible. Nuclear speckles and Cajal bodies, while distinct in composition, frequently share interfaces that facilitate exchange^25^. In the cytoplasm, P-bodies and stress granules often dock against one another, forming composite droplets with distinguishable biochemical identities and an identifiable interface^26,27^. These condensates also dynamically adjust their internal structure, rapidly assembling, disassembling, or reorganizing in response to cellular stress, metabolic changes, or signaling events^28–30^. Stresses like heat shock or transcriptional inhibition can disrupt nucleolar substructure, promoting partial mixing of phases, while recovery re-establishes internal segregation^29,31^.

Despite these striking observations, we do not completely understand how cells encode multiphase architectures at the molecular level. What sequence features determine whether macromolecules co-partition into a single mixed phase or segregate into two, three, or more distinct subdomains within the same condensate? What determines the extent of partitioning? How do cells encode and dynamically regulate this miscibility in response to environmental signals? These fundamental questions about the molecular grammar of multiphase organization remain largely unanswered.

In vitro studies have shown that proteins with distinct disordered regions can form nested condensates^32^, RNAs can drive sub-compartmentalization^33,34^, and synthetic polymers with different chemical properties demix when co-condensed^35–40^. These examples illustrate that multiphase architectures can arise from minimal components, yet the governing principles remain largely descriptive. A key challenge is to move beyond phenomenology and develop quantitative frameworks that link molecular-scale interaction parameters to emergent multiphase behavior.

Here, we show that sequence-encoded molecular interactions can precisely program the internal architecture of multiphase condensates. We propose that the key to this control is an emergent property of the condensed state itself. Associative phase separation, such as complex coacervation^41,42^, can drive macromolecules above the overlap concentration, shifting the system into a semi-dilute regime where polymer chains interpenetrate and inter-molecular contacts are frequent. In this emergent state, subtle molecular preferences, which are largely irrelevant in dilute solution, become dominant organizers of internal architecture. Using a programmable DNA system^43^, we show that when multiple species co-condense, even modest differences in homotypic interactions, of just a few kT, can overcome the entropic cost of demixing, driving spontaneous spatial segregation without membranes or scaffolds. Critically, we achieve quantitative control: partitioning between phases scales precisely with interaction energy, and using orthogonal interaction domains we generate condensates with up to four coexisting phases. A modified sticker-spacer model^44^ accurately predicts these behaviors. These principles extend to RNA-peptide systems, establishing general design rules for sequence-programmed organization. Together, these findings establish phase separation as more than a concentration mechanism: within condensates, weak molecular preferences become powerful organizational forces that drive hierarchical self-assembly.

## Results

### Hybridization-mediated cross-links drive spontaneous sub-compartmentalization in DNA condensates

We previously showed that the tetravalent cation spermine induces associative phase separation of single-stranded DNA^43^. This process concentrates DNA above its critical overlap concentration, placing it in the semi-dilute regime. Crucially, within this condensed phase, DNA strands retain their ability to engage in sequence-specific hybridization^43^. To test whether differential molecular interactions drive internal phase separation, we designed two single-stranded DNA oligonucleotides: a 90-nucleotide homopolymeric polyT (T90), and a corresponding patchyDNA variant containing four evenly spaced palindromic GGCC motifs separated by poly(T) stretches (**Figure 1A**). These GGCC “stickers” mediate weak, reversible intermolecular hybridization, while the intervening poly(T) regions act as inert non-hybridizing spacers. Adjacent stickers are separated by ≥15 T bases (**Table S1** and **Supp. Note 1**) in order to minimize cooperativity in hybridization between adjacent sites^45,46^. By design, both oligonucleotides are 90 nucleotides long and have identical charge but differ only in their capacity to form base-pairing-mediated cross- links, allowing us to isolate the contribution of specific molecular interactions to condensate behavior.

**Figure 1:**
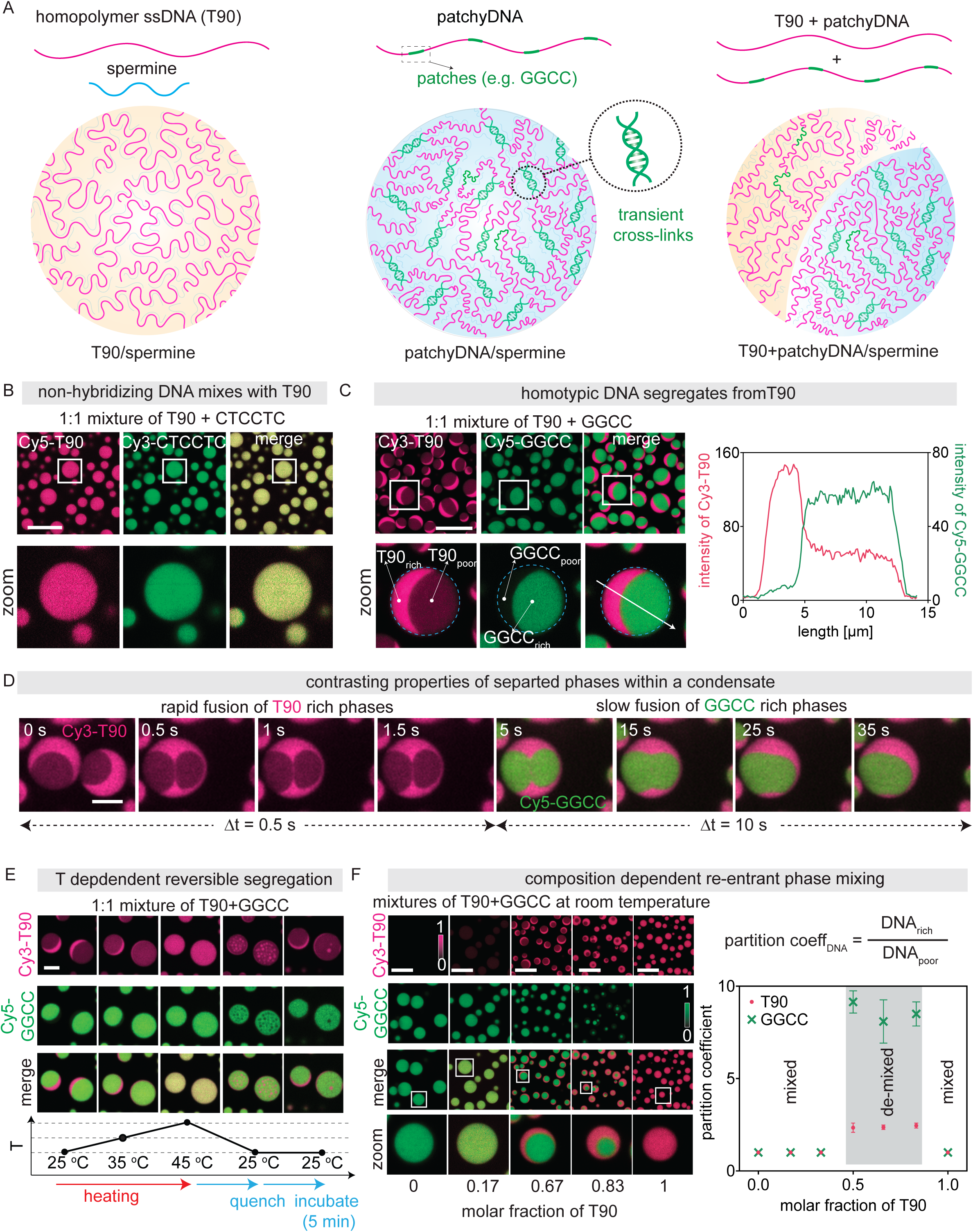
DNA condensates exhibit composition- and temperature-tunable internal demixing: **(A)** Schematic. T-90 forms electrostatically driven condensates with the polycation spermine, while patchyDNA strands carry short hybridization patches (e.g., GGCC) that create transient, sequence-specific cross-links. Co-condensing T-90 with patchyDNA results in spontaneous segregation of the two species within DNA/spermine condensates, enabling programmable sub-compartmentalization. (B) Representative fluorescence micrographs showing that non-hybridizing patchyDNA (CTCCTC patches) mixes with T-90. (C) Representative fluorescence micrographs (left) and corresponding intensity traces (right) showing that T-90 chains segregated from patchyDNA (GGCC patches) and formed bi-phase condensates with a T-90 rich and a patchyDNA rich phase. (D) Fluorescent micrographs showing coalescence of two T- 90+GGCC condensates. T90-rich domains fuse rapidly, while GGCC-rich domains fuse slowly, indicating distinct viscoelastic properties for the coexisting internal phases. (E) Representative fluorescence micrographs showing reversible mixing and de-mixing of the two DNA species within T90+GGCC condensates with temperature (T). (F) Representative fluorescence micrographs showing T90+GGCC condensates (left), and corresponding partition coefficients (right) at indicated mole fraction of T-90. All DNA/spermine condensates (B-F) were generated at the following conditions: Total DNA = 10 µM, spermine = 4 mM in 10 mM Tris pH 7.0. T = 22°C in (F). Scale bars in (B), (C), and (F): 10 µm, in (D): 5 µm, in (E): 3 µm.

The strength of individual hybridization-mediated cross-links is captured by the dimensionless parameter ε = -ΔG/RT, where ΔG is the theoretical free energy associated with dissociation of a single pair of patches and R is the gas constant (**Table S1, Supp. Note 1**)^47,48^. PatchyDNA with GGCC patches had a predicted ε = 12.5, in 1 M salt at 22 °C, although the effective ε will depend on the precise ionic environment in the condensed phase^49^. In contrast, T-90 does not have an associative patch and thus is not expected to form hybridization mediated cross-links (ε = 0).

We mixed the T-90 and the patchyDNA with GGCC patches at equimolar ratio (5 µM each in 10 mM Tris, pH 7.0) and induced complex coacervation by adding 4 mM spermine. The reaction was doped with fluorescently labeled DNAs (40 nM each of Cy3 labeled T90 and Cy5 labeled patchyDNA, respectively) in order to facilitate visualization by fluorescence microscopy. Addition of spermine triggered complex coacervation^50,51^ as evidenced by increased solution turbidity and the appearance of numerous spherical droplets (**Figure 1B-C**). Within droplets, however, T-90 and GGCC DNAs exhibited striking spontaneous spatial segregation, forming two coexisting DNA-rich phases: a T90-enriched phase and a GGCC-enriched phase (**Figure 1C**). These coexisting phases were liquid like and shared a smooth, continuous internal interface. When two or more droplets came in proximity, they coalesced to form a larger droplet that relaxed to a spherical shape (**Figure 1D and Supp. Movie 1**). Upon fusion, the like-domains coalesced: T-90-rich regions merged into a single, unified compartment, as did the GGCC-enriched regions (**Figure 1D and Supp. Movie 1**). The two phases, however, remained segregated throughout this process, thus preserving the internal biphasic architecture (**Figure S1**). In contrast, similar experiments performed with a control patchyDNA which cannot self-hybridize, (patch sequence CTCCTC), produced homogeneously mixed droplets with T-90 with no evidence of internal phase separation (**Figure 1B**).

To characterize the stability of this multiphase organization, we first quantified the degree of molecular partitioning. We measured the partition coefficient, K, defined as the ratio of the concentration of a DNA component in its preferred (enriched) phase to that in the non-preferred (depleted) phase (see partition coefficient section in **Methods** and **Figure S1**)^52^. PatchyDNA with patch sequence GGCC exhibited strong enrichment (K_GGCC_ = 9.1 ± 0.6, mean ± SD, n = 26 droplets), while T90 showed more modest but reproducible partitioning (K_T-90_ = 2.34 ±0 .25, mean ± SD, n = 181) (**Figure S1C**). Crucially, these values were remarkably uniform across hundreds of droplets within a sample and reproducible across independent experiments (**Figure S1**). To confirm that this highly consistent organization represents a thermodynamic endpoint rather than a kinetically trapped state, we mixed pre-formed T90-only and GGCC patchyDNA-only condensates. These pre-formed droplets readily fused and reorganized into biphasic structures indistinguishable from those formed by pre-mixing DNAs before spermine addition (**Figure S1A**).

The resulting partition coefficients were comparable across both preparation methods (**Figure S1C**), and remained stable over time (**Figure S1B**), suggesting that the observed multiphase architecture is a robust, consistent, and pathway-independent equilibrium state.

We next examined how this phase behavior responds to external perturbations. Base-pairing interactions are particularly temperature sensitive^47^. As we increased the temperature, the T-90- rich and GGCC-rich phases became increasingly miscible. The partition coefficient of T-90 (K_T90_) progressively decreased from 2.4 ± 0.2 (mean ± SD, n = 62) at 25°C to unity at 45°C (**Figure 1E, Figure S2 and Supp. Movie 2**), corresponding to complete mixing of the two DNA species into a single homogenous phase. This process was fully reversible: upon cooling, the biphasic architecture re-emerged, restoring the initial partitioning (**Figure 1E and Supp. Movie 2**). The reversibility of temperature-induced mixing further indicated that phase segregation arises from an underlying equilibrium process.

Next, we examined how internal organization depends on component stoichiometry: another thermodynamic variable that, like temperature, can modulate phase coexistence^13,52^. To map the compositional phase diagram, we systematically varied the molar ratio of T-90 and GGCC patchyDNA while maintaining a constant total DNA concentration (10 µM). Biphasic condensates emerged only within a defined compositional window (when molar fraction of T90 was between 0.5 and 0.83) (**Figure 1F**). Outside this range, the system formed a single, homogeneously mixed DNA-rich phase. Within the two-phase regime, the concentration of each DNA in its respective enriched phase remained constant, and only the relative phase volumes changed with input ratio (**Figure 1F**). This lever-rule behavior is a hallmark of equilibrium phase separation^53,54^ and confirms that the observed demixing reflects coexistence between two thermodynamically stable states with fixed compositions. The width of the miscibility gap likely reflects a balance between the enthalpic gain from homotypic GGCC base-pairing interactions and the entropic cost of demixing, as well as any residual heterotypic interactions that oppose complete segregation^55,56^. Altogether, these results demonstrate that hybridization-mediated cross-linking drives spontaneous spatial segregation of DNAs within condensates, and that the resulting internal organization reflects a thermodynamically stable state.

### Coexisting phases retain distinct material properties encoded by sequence-specific crosslinks

We next asked whether the spatially segregated phases create two distinct microenvironments within the same droplet. Our previous work established that hybridization-mediated crosslinks dramatically alter condensate material properties^43^. To determine if these contrasting properties are preserved within biphasic condensates, we measured molecular diffusivity, viscosity, and phase stability within each domain. Fluorescence recovery after photobleaching (FRAP)^57^ revealed that DNA mobility within each phase closely matched that of single-component condensates (**Figure 2A-C**). T90 molecules in the T90-rich phase diffused rapidly (D _self_ = 0.17±0.02 µm²/s, mean ± SD, n = 3), comparable to pure T90 condensates (**Figure 2B-C**). In contrast, GGCC-patchyDNA in its enriched phase showed 10-fold slower diffusion (D = 0.017±0.001 µm²/s mean ± SD, n = 5), consistent with the formation of a cross-linked network through base-pairing interactions. Minority components exhibited asymmetric behavior: T-90 maintained high diffusivity even within the viscous GGCC-rich phase (**Figure S2C**). In contrast, the GGCC patchyDNA in the T-90-rich phase diffused as rapidly as T-90 itself (D_self_ = 0.12 ± 0.02 µm^2^/s, mean ± SD, n = 5 **Figure S2D**). This asymmetry likely arises because crosslinking requires sufficient polymer density. At the dilute GGCC concentrations in the T90-rich phase (∼10% of the concentration in the pure GGCC patchy-DNA condensates), molecules would rarely encounter hybridization partners and thus behave similar to non-interacting chains.

**Figure 2:**
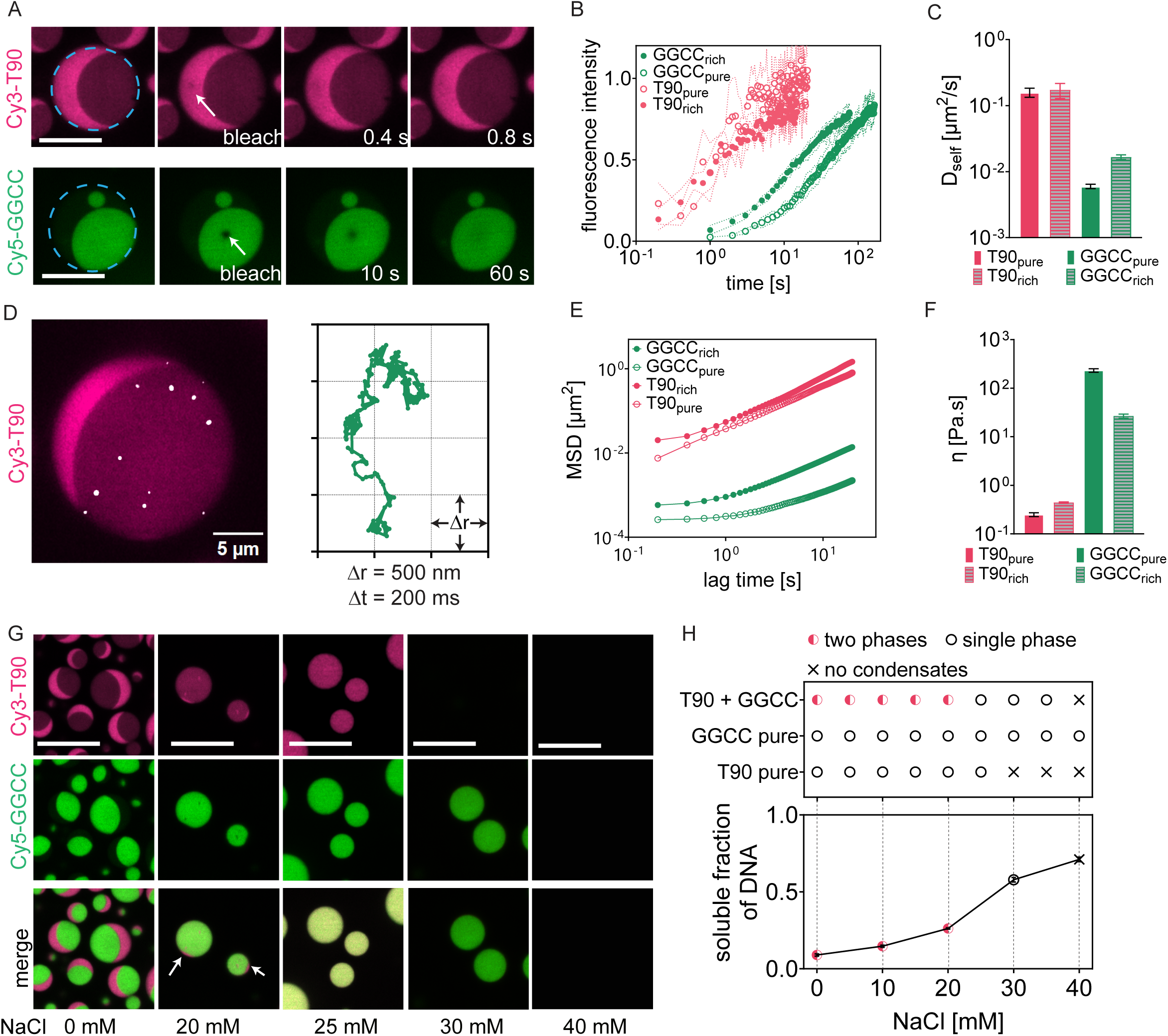
Sequence-dependent material and phase properties of biphase condensates: (A) Representative micrographs showing FRAP of T90 (top) and GGCC (bottom) within their respective enriched domains in T90+GGCC condensates. (B-C) Normalized recovery curves (B) and corresponding self-diffusivity (D_self_) (C) indicate that the non-hybridizing T90 chains were highly diffusive in T90-rich domains of T90+GGCC condensates (T90_rich_), with a mobility similar to that in pure T90 coacervates (T90_pure_). In contrast, transient, hybridization-mediated cross- linking slows the diffusion of GGCC chains in both their pure condensates (GGCC_pure_) and in GGCC rich phases in T90+GGCC condensates (GGCC_rich_). (D) Micrograph showing embedded fluorescent beads (radius, 50 nm) in a T90+GGCC condensate (left) and representative trajectory of a bead in the T90-rich phase (right). (E-F) Mean squared displacement (MSD) of embedded beads (E) and corresponding viscosity (F) showing that T90 rich phase had low viscosity, comparable to T90-only coacervates. In contrast, GGCC-rich phase was ∼100 times more viscous than T90 rich phase and resembled pure GGCC condensates. (G) Representative micrographs showing that GGCC-rich phase exhibits higher resistance against NaCl mediated dissolution, compared to T90-rich phase. (H) Phase diagram showing stability of indicated DNA/spermine condensates (top) and soluble DNA fraction for T90+GGCC condensates (bottom) as a function of NaCl concentration. Scale bars: (A):10 µm, (D): 5 µm, and (G): 12 µm.

To determine whether these mobility differences are reflected in broader mechanical properties, we measured viscosity in each phase using particle tracking microrheology^58^. We embedded 50 nm fluorescent beads and tracked their motion to extract mean squared displacement (MSD) over time (**Figure 2D**). For purely viscous fluids, the MSD increases linearly with lag time (t) following the relation: MSD = 4D×t + NF, where D is the diffusivity of the bead and NF is the noise floor. For both phases, MSD increased linearly with lag time, consistent with purely viscous behavior (**Figure 2E**). The viscosity (η) of each phase was calculated using the Stokes-Einstein relation. The measured viscosities revealed a striking contrast: the T90-rich phase exhibited low viscosity (η_T-90-rich_ = 0.44 ± 0.01 Pa.s, mean ± SD, n = 7) whereas the GGCC-rich phase was substantially more viscous (η_GGCC-rich_ = 26 ± 2 Pa.s, mean ± SD, n = 6), representing a nearly 60- fold difference in material properties across the phase boundary (**Figure 2E-F**). These values closely matched those of the corresponding single-component condensates, confirming that sequence-specific hybridization still governs the network connectivity within the demixed phases (**Figure 2F**).

We next asked whether the two coexisting phases differ in their resistance to salt-induced dissolution, as a readout of phase stability^51,59^. In complex coacervates, increasing concentrations of monovalent salt typically disrupt phase integrity by screening electrostatic interactions between polyanions and polycations^43,51,59^. We gradually titrated NaCl into biphasic droplets and tracked the dissolution of each phase. The T-90-rich phase dissolved at a critical salt concentration (C*_T90- rich_) of 30 mM, similar to pure T-90 condensates (**Figure 2G**). In contrast, the GGCC-rich phase persisted until C*_GGCC-rich_ = 40 mM NaCl, matching the enhanced stability of pure GGCC-only condensates (**Figure 2H**). This difference in critical salt concentration reflects the additive contribution of reversible base-pairing to phase stabilization^43,60^. The sequential loss of phase integrity with increasing salt confirms that each domain retains a distinct molecular interaction network, despite coexisting within a continuous droplet. Together, these results demonstrate that complex internal organization can arise spontaneously from simple molecular rules: differential hybridization alone is sufficient to generate coexisting, stable phases. Each domain retains the material properties encoded by its dominant molecular component, allowing a single droplet to contain subregions that differ by orders of magnitude in viscosity and mobility, yet share a smooth, continuous interface.

### A critical interaction energy triggers multiphase organization

Having established that sequence-specific interactions drive phase separation within DNA coacervates, we next asked whether this behavior could be systematically tuned by modulating the base-pairing strength of the interacting patches. To test this, we designed a panel of patchyDNAs with interaction energy, ε, spanning values from ≈5 to ≈20 (calculated at 1 M NaCl and 22°C, **Figure 3A**, see **Methods and Supplementary Note 1**). Each patchyDNA was mixed with equimolar amounts of the non-binding T-90 DNA (ε = 0) and subjected to complex coacervation with spermine (**Figure 3A-B**). These experiments revealed a sharp transition in phase behavior (**Figure 3A-B**). Sequences with weak hybridization (ε ≤ 8.4) formed homogeneous, single-phase condensates, while those with stronger interactions (ε ≥ 12.5) spontaneously demixed into biphasic structures with T-90-enriched and patchyDNA-enriched domains (**Figure 3A-B and Figure S3**). This critical threshold, ε_critical_ (8.4 < ε_critical_ < 12.5), demarcates complete miscibility from phase separation, demonstrating that a minimum interaction asymmetry is required to overcome the mixing entropy^53,55^.

**Figure 3:**
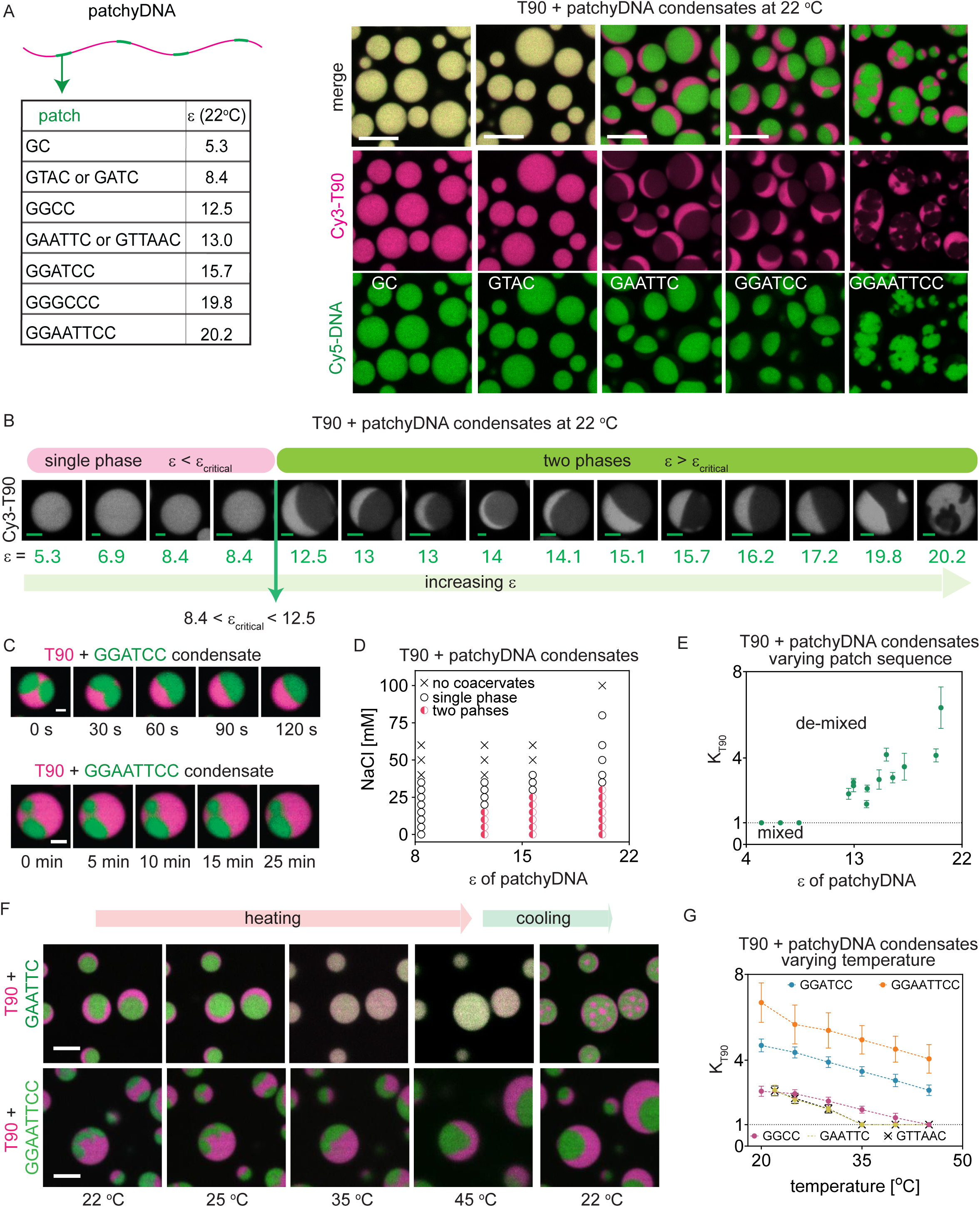
A critical hybridization energy triggers multiphase organization: (A-left) Representative patch sequences and their corresponding hybridization energies (ε) at 22° C. (A- Right) Representative fluorescent micrograph of T90+patchyDNA/spermine condensates showing that patchyDNA with weak patches (GC and GTAC) mix completely with T90 and form single phase condensates. PatchyDNA with relatively strong patches (GAATTC, GGATCC and GGAATTCC) segregate from T90 chains, forming condensates with a T90 rich and a patchyDNA rich phase. (B) Representative fluorescent micrographs of T90+patchyDNA condensates for 15 distinct patchyDNAs with indicated ε at 22°C showing that a critical ε is required to form bi-phase condensates. (C) Representative fluorescent micrographs showing fusion of GGATCC-rich (top) and GGAATTCC-rich (bottom) phases within their corresponding T90+patchyDNA condensates. Phase diagram of T90+patchyDNA condensates with NaCl as a function of ε for patchyDNA. Partition coefficient of T90 (K_T90_) in bi-phase condensates as a function of ε. K_T90_ = 1 indicates T90-rich and patchyDNA-rich phases are completely mixed. (F) Representative fluorescent micrographs showing bi-phase condensates of T90+GAATTC reversibly switch between mixed and demixed internal organization with temperature (top), whereas T90+GGAATTCC condensates (bottom) remain demixed even at 45°C. (G) Partition coefficient of T90, K_T90_, in bi- phase T90+patchyDNA condensates with indicated patch sequences as a function of temperature. Scale bars in A, B, C and F represent 10 µm, 2 µm, 2 µm and 3 µm, respectively.

Across all biphasic condensates, each phase preserved the material features of its dominant component resulting in two phases with distinct and highly tunable material properties (**Figure S4**). The T-90-rich phase (consisting of ≈70% T-90 DNA, **Figure S4**) consistently exhibited low viscosity, high diffusivity, and low salt stability, comparable to pure T-90 condensates, regardless of which patchyDNA variant occupied the adjacent phase (**Figure S4**). In contrast, the patchyDNA-rich phases displayed material properties that were systematically programmed by hybridization energy (ε) (**Figure S4**)^43^. These differences were vividly illustrated by the droplet coalescence dynamics (**Figure 3C and Supp. Movie 3, 4**). Similar to our observation with the T- 90+GGCC/spermine system (**Figure 1D**), all biphasic droplets exhibited phase-specific coalescence: when two droplets fused, their respective T-90 and patchyDNA-rich domains selectively merged with their counterparts (**Figure 3C and Figure S4**). While this selective fusion behavior was universal, the dynamics were starkly asymmetric. The T-90-rich domains consistently coalesced within seconds (**Figure S4E-F**). In contrast, the fusion timescale for the patchyDNA domains was exquisitely sensitive to ε: approximately 2 µm GGATCC-rich domains (ε = 15.7) fused within 2 minutes, while GGAATTCC-rich domains (ε = 22.9) of comparable size remained only partially fused even after 25 minutes (**Figure 3C**).

These macroscopic dynamics were underpinned by sequence-encoded changes in mobility at the molecular scale^43^. As ε increased, the molecular diffusivity within the patchyDNA-rich phase decreased exponentially (**Figure S4G-H**), while its resistance to salt-induced dissolution systematically increased (**Figure 3D and Figure S5**). Crucially, this exponential scaling of diffusivity with ε quantitatively reproduced the behavior we previously established for pure, single-component patchyDNA condensates^43^. Thus, despite being embedded within the same droplet, the two coexisting phases could differ by over 100-fold in viscosity and molecular mobility. This systematic scaling confirms that sequence-encoded interaction energy quantitatively governs both the onset of phase separation and the material properties of the resulting DNA-rich phases.

### A unified scaling law governs demixing

To quantify the mixing-demixing transition, we examined how T-90 partitions between two phases as a function of hybridization energy, ε (**Figure 4A-C**). We focused specifically on T-90 partitioning because its input concentration and labeling efficiency could be reliably standardized across samples; in contrast, variations in fluorophore labeling or minor inaccuracies in stock concentrations between the various patchyDNAs could potentially introduce biases in measurements of their relative enrichment. Interestingly, we observed that as ε increased, T-90 was progressively excluded from the patchyDNA-rich phase (**Figure 3B**): its concentration decreased within the patchyDNA-rich domain while modestly increasing in the T-90-rich domain. To quantify this partitioning, we calculated the partition coefficient K_T-90_, defined as the ratio of T-90 concentration between phases (**Figure 3E**). Plotting K_T-90_ against ε for our panel of sequences revealed a sharp, two-regime behavior: the system remained fully mixed (K_T-90_ = 1) until a critical threshold (ε_critical_), after which partitioning increased steadily with interaction energy (**Figure 3E**). Notably, we observed a nearly linear relationship between K_T-90_ and ε in this second regime, indicating that phase segregation is directly correlated to interaction strength (**Figure 3E and Figure S3C**).

**Figure 4:**
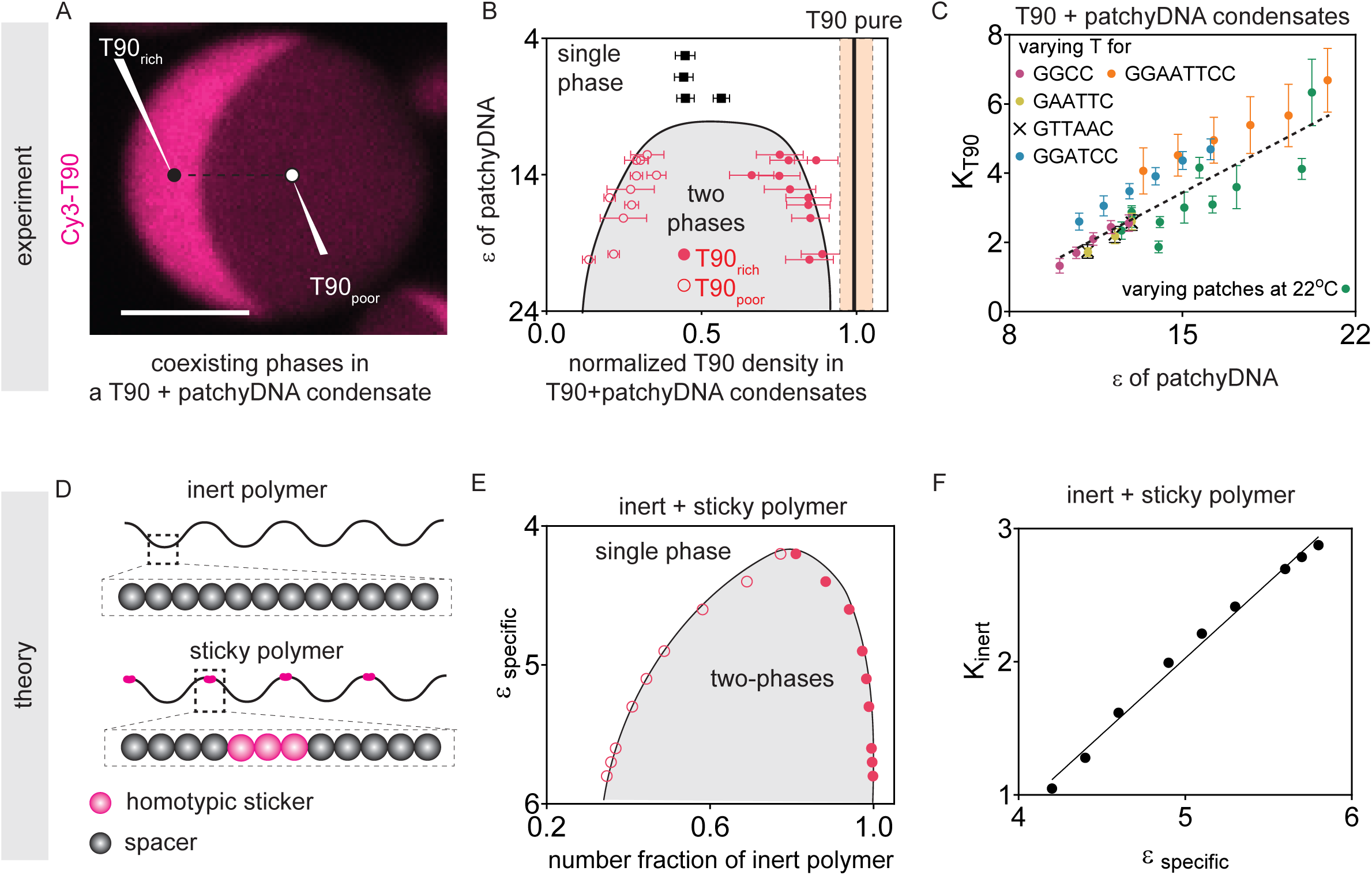
Universal scaling of internal demixing with sticker hybridization energy from experiments and sticker-spacer theory: (A) Representative fluorescence micrograph showing T90_rich_ and T90_poor_ compartments within a T90+patchyDNA condensate. (B) Phase diagram of T90+patchyDNA condensates showing the normalized T90 density in single phase (square) and two-phase (internally demixed) condensates (circles). The T90 density was normalized to 1 (black line) corresponding to the concentration in pure T90 coacervates with shaded band indicating SD (n= 150). (C) Master plot showing that the partition function of T90 (K_T90_) in T90+patchyDNA condensates increased linearly with ε, whether it was varied by changing the patch sequence or by temperature. The black-dotted line is the linear fit to the experimental data with a slope, 0.38 ± 0.01 (mean ± 95% CI). (D) Schematic of the theory model. An inert polymer consists of 88 identical inert monomers. In contrast, the sticky polymer, also containing 88 monomers, was designed with 4 homotypic stickers, separated by 19 spacer monomers, creating a regular periodic structure. (E) Theory, with same analysis as in B. Calculated phase diagram for binary mixtures of the inert polymer with sticky polymers, as a function of sticker binding affinity. (F) Theory, with same analysis as in C. Calculated partitioning of the inert polymer versus sticker binding affinity (slope, 1.1 ± 0.1 (mean ± 95% CI)). Scale bar in A: 5 µm.

To test the universality of our findings, we next modulated interaction energy orthogonally, by tuning temperature for fixed DNA sequences. Because hybridization free energy is temperature-dependent, it allows us to tune ε without altering the molecular components (**Supplementary Table S2**)^47,48^. We measured K_T-90_ for various patchyDNA sequences across a range of temperatures from 25°C to 45°C (**Figure 3F-G**). As the temperature increased, we observed progressively reduced partitioning of T-90 into the patchyDNA-rich phase, as expected (**Figure 3F-G**, **Figure S6**). Across these varied conditions, measured K_T-90_ consistently displayed a linear dependence on calculated ε (**Figure S6C**). Remarkably, whether ε was tuned by sequence or temperature, all measurements collapsed onto a single K_T-90_-ε line (slope = 0.38±0.01; 95% CI, **Figure 4A-C**). This universal scaling behavior, independent of the mechanism by which ε is tuned, demonstrates that hybridization energy is the primary driver for the extent of demixing. Once ε exceeds a critical value, the enthalpic gain from base-pairing interactions overcomes the entropy of mixing, driving phase separation.

### Associative polymer theory captures sequence-programmed phase separation

The near-linear relationship between molecular interaction energy and macroscopic phase segregation motivated us to develop a theoretical framework that could explain these phenomena from first principles. The observed separation of patchyDNA and T-90 is intuitive from the perspective of Flory-Huggins theory, which shows that even slightly different polymers will phase separate if they are long enough^53,54,61^. However, there are two difficulties in capturing the details of our system within the mean-field χ parameter of the Flory-Huggins model^53,55,62^. First, our DNA strands have fixed valence: each strand contains a defined number of patches that can mediate hybridization between two strands, constraining the connectivity of the network (**Figure 1A**)^43^. This differs from mean-field models that assume uniform interactions along the polymer. Second, the lattice approximation of Flory-Huggins implicitly introduces a length scale corresponding to the interaction scale. However, our system contains two distinct interaction scales: non-specific electrostatic attractions mediated by spermine and sequence-specific hybridization between associative patches^43^. Both of these shortcomings are addressed by the “associative polymer” model first introduced by Semenov and Rubinstein^44,63–65^, which explicitly accounts for the statistics of reversible cross-linking with fixed valence (see **theory SI** for details). This model contains three key contributions to the free energy. The mixing entropy of the polymers follows the standard form:

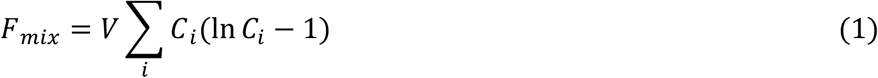

where *C_i_* is the concentration of each polymer species and *V* is volume. The next term is the binding energy and entropy of patch-patch interactions

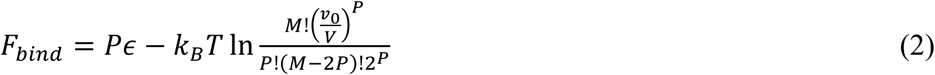

where *M* is the total number of stickers in the system, *P* is the number of patch-patch bonds, and *v_0_* is the effective patch search volume^44,66^. Finally, Semenov and Rubinstein accounted for all interactions not involving the patches via a third order virial expansion. The bonding term *F_bond_* favors locally high concentrations of patches, which increases both the combinatorial entropy and energy of bond formation. This driving force for patch concentration is opposed by both the mixing entropy and repulsive interactions captured by the virial terms.

To apply this framework to our system, we must account for the spermine-mediated interactions that drive initial condensation. While models of low valence crosslinkers are available^67^, the unknown partitioning of spermine between the dense and dilute phases along with its presence in both dense phases complicates direct application. We therefore employ a perturbative approach in which we define *C_0_* as the optimal polymer concentration in a fluid lacking patch-patch interactions (i.e., pure T-90 condensates). To lowest order, these deviations can be accounted for with a quadratic potential,

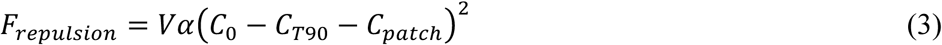

where α is a constant. Introducing patchyDNAs will disrupt the T-90 fluid in two ways. First, the attractive patches will try to maximize their concentration to minimize *F_bond_*. However, the increased persistence length of the hybridized patches, relative to single stranded DNA, will locally swell the network. The quadratic potential of Eq. 3 captures the total disruption of the T-90 fluid due to sub-optimal spermine density, electrostatic repulsion, and steric interactions.

The total free energy is given by the sum of Eqs. 1-3, which can be expressed in terms of a composition parameter = 𝐶_*patch*_/𝐶_*T90*_ (see **theory SI**). We used a pseudo-two phase approximation, neglecting the dilute phase, to determine when the condensate separates into two distinct phases using a common tangent construction^68^. The resulting phase diagram successfully captures the key experimental observations, predicting a critical patch affinity for separation of the T-90 and patchyDNA molecules and increased partitioning with increasing interaction strength (**Figure 4D-F, Figure S7**). The model provides a clear physical mechanism for this behavior: below the critical hybridization energy, the entropy of mixing dominates, resulting in a single homogeneous phase. Above this threshold, the patchy DNAs preferentially associate into clusters. These clusters exclude T-90 which segregate through non-specific electrostatic interactions. The excluded volume repulsion between these distinct networks, one stabilized by base-pairing, the other by electrostatic crosslinks, helps separate the phases to create an immiscible two-phase condensate.

While the model captures these qualitative features, we note two quantitative discrepancies. First, the magnitude of the critical affinity is different between the theory and experiment, which we attribute to the difference between the calculated ε at 1 M salt and the altered environment in the condensed phase. Second, the experimental phase diagram (**Figure 4B**) is somewhat more symmetric than the calculated phase diagram (**Figure 4E**), which is likely due to the theory’s inability to capture swelling due to the reduced persistence length of hybridized DNA. Despite these offsets, our model provides a clear design principle: orthogonal interactions of sufficient strength should drive segregation into a corresponding number of phases. We next sought to test this prediction directly.

### Programming condensates with multiple coexisting phases

Our theoretical framework predicts that phase separation depends only on pairwise interaction differences, implying that *n* orthogonal sequences should yield *n* coexisting phases, provided each pairwise interaction strength exceeds ε_critical_. To test this hypothesis, we designed orthogonal patchyDNA sequences with minimal cross-hybridization propensity (patch sequences GGATCC, GGGCCC; see **Table S3**). When T-90 was mixed with these patchyDNAs in equimolar amounts and subjected to spermine-induced coacervation, we observed DNA-rich droplets that spontaneously organized into three distinct spatial domains, each enriched in a single DNA species (**Figure 5A**). These co-existing phases formed smooth and stable interfaces, similar to our observation with two-phase condensates.

**Figure 5:**
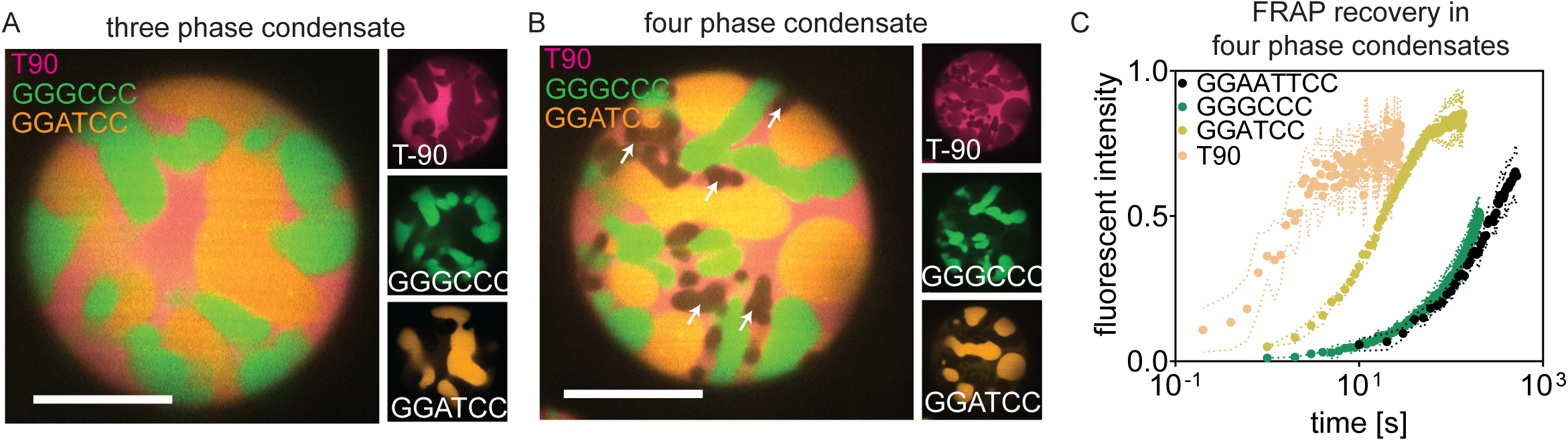
Programming condensates with multiple coexisting phases: (A) Left: Representative micrographs of a three phase condensate from a 1:1:1 molar mixture of T90 and two patchyDNA species bearing GGATCC and GGGCCC stickers. Right: Single-channel views highlighting the T90-rich, GGATCC-rich, and GGGCCC-rich subdomains within the same droplet. (B) Similar to A, but for a four-phase condensate assembled from T90 and three patchyDNAs with following patches: GGATCC, GGGCCC, GGAATTCC. The GGAATTCC dense phase is indicated by white arrows. (C) FRAP recovery of the indicated DNA in their respective rich compartments within a four-phase condensate demonstrating domain-specific mobilities. Scale bars: 10 µm.

Building on these results, we next introduced a third orthogonal patchyDNA (patch sequence GGAATTCC). Combining T-90 with all three patchyDNA variants produced droplets with four distinct, stable phases clearly resolved by multicolor fluorescence imaging (**Figure 5B**). Notably, no templates or external cues were required, and this internal architecture emerged purely from the encoded molecular interactions. Quantitative characterization revealed that each phase maintained distinct material properties. FRAP measurements showed that molecular diffusivity within each domain matched predictions based on its dominant component (**Figure 5C and Figure S8**). The T90-rich phase exhibited consistently high molecular mobility, while the patchyDNA- rich phases displayed progressively slower dynamics scaling with their respective hybridization energies (**Figure 5C and Figure S8**), similar to our earlier observations. In principle, this approach could support even greater phase complexity limited mainly by the challenge of designing a set of truly orthogonal DNA interaction domains. Nevertheless, our results reveal general design principles for engineering multiphase condensates, encoding diverse molecular and mechanical environments within a single, continuous droplet.

### Sequence-encoded multiphase RNA condensates

Many cellular condensates are ribonucleoprotein assemblies where RNA structure and interactions play organizing roles. To test whether our design rules extend beyond DNA, we examined RNA-peptide condensates, as models for RNP granules^33,50,69,70^. We designed a two-component RNA system: an unstructured 60 nucleotide polyU sequence (rU-60) and a patchyRNA variant containing four evenly spaced self-complementary rGGCC motifs within a polyU backbone (**Table S4**)^43^.

Mixing these RNAs with 10-mer oligo-L-lysine (K10) produced coacervate droplets (**Figure 6A**). We doped a small amount of Cy3 labeled U-60 (<1% of total rU-60) to facilitate visualization by fluorescence microscopy. As with DNA, the internal organization of the droplets depended on stoichiometry of the RNA components. When either component was in excess, the RNA chains remained mixed and formed single phase condensates (**Figure 6A**). Spontaneous demixing occurred within a specific compositional window, when the molar fraction of rU60 was between 0.66 and approximately 0.38, yielding distinct rU60-enriched and patchyRNA-enriched domains with shared interface (**Figure 6A-B**).

**Figure 6:**
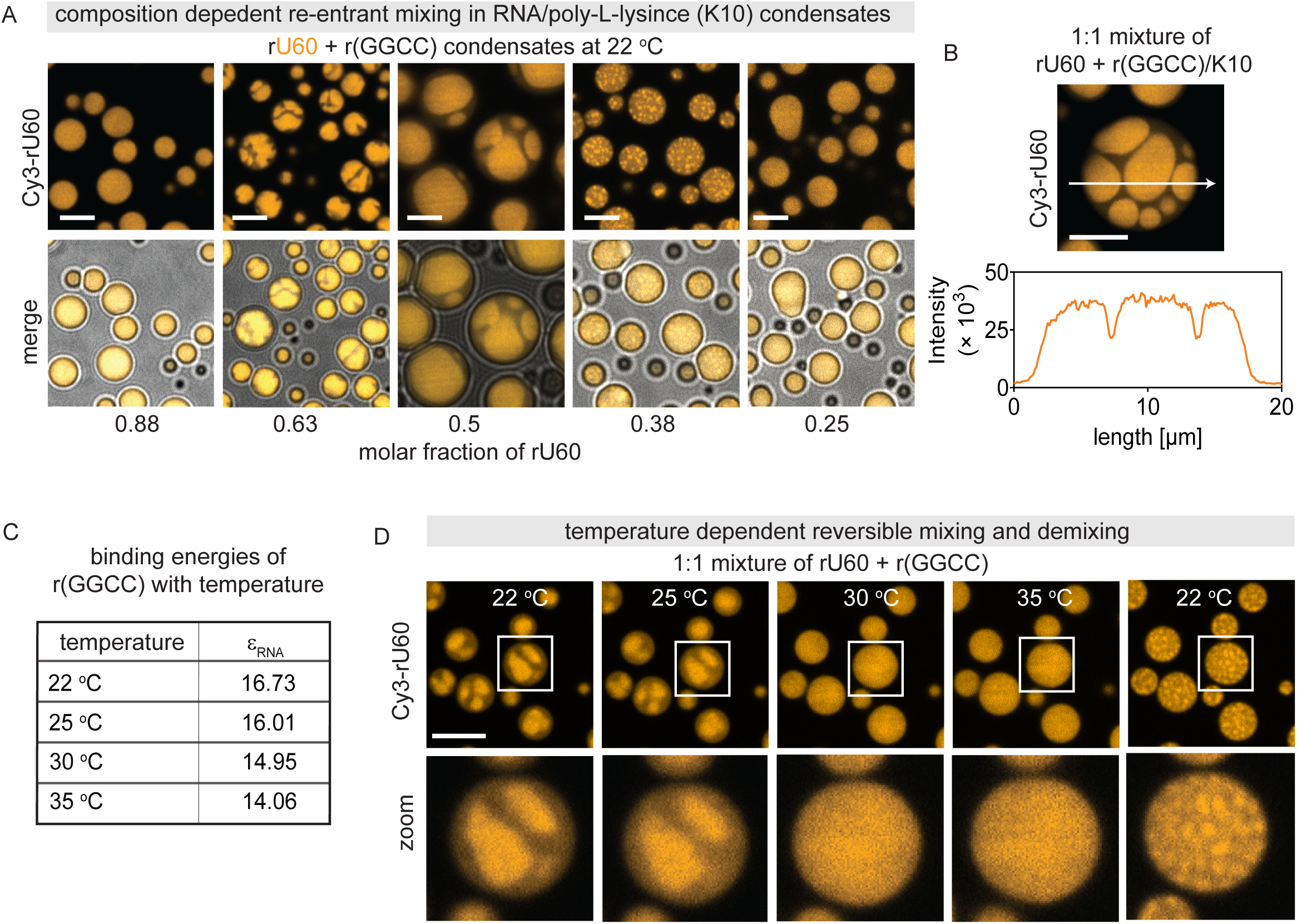
Sequence-encoded multiphase RNA condensates: (A) Composition-dependent phase behavior in RNA/poly-L-lysine (K10) condensates at 22°C formed from 60-nt rU60 and a 60-nt patchyRNA carrying four GGCC stickers. Top: fluorescence micrographs; bottom: corresponding fluorescence-brightfield merges. Varying the molar fraction of rU60 yields re-entrant mixing: two- phase demixing at intermediate compositions and single-phase mixing toward rU60-rich or r(GGCC)-rich limits. (B) Representative fluorescent micrograph (top) and a corresponding intensity line scan across the droplet (bottom) a two-phase condensate formed in a 1:1 mixture of rU60 + r(GGCC). (C) ε of r(GGCC) patches at indicated temperatures. (D) Representative fluorescence micrographs showing reversible dissolution and formation of two-phase RNA condensates with temperature. Scale bars in A: 5 µm, and D: 4 µm.

The temperature response of these RNA condensates also paralleled the DNA system, highlighting a shared thermodynamic basis (**Figure 6C-D, Table S5**). Heating biphasic condensates from 22°C to 30°C progressively mixed the phases; at 30 °C the condensates were fully homogeneous, indicating that a critical hybridization interaction is required for demixing (**Figure 6C-D and Figure S9**). Further increasing the temperature to 35°C produced no additional morphological change (**Figure 6D and Figure S9**), confirming the system had reached a stable, single-phase state. Importantly, this transition was fully reversible: upon cooling, numerous sub-micrometer rU60-rich domains nucleated (**Figure 6D**). These internal U60-rich domains coalesced to re-establish the equilibrium two-phase architecture within minutes (**Figure S10**).

Although the phase transition was fully reversible, we discovered that the droplets could exhibit striking structural memory of their previous state^71^ when subjected to brief thermal pulses (**Figure S10**). A brief 1 min thermal pulse at 30°C appeared to fully mix the phases as assessed by fluorescence microscopy, yet upon cooling, new phases preferentially re-formed at their prior locations (**Figure S10A-B**). In contrast, extended incubation (10 min at 30°C) successfully erased this memory, leading to random nucleation of rU-60-rich islands throughout the droplet (**Figure S10C**). These observations suggest that residual molecular-scale organization can persist over short timescales despite the macroscopically uniform appearance, and template the location of subsequent phase separation^72^.

## Discussion

Our work establishes that sequence-encoded molecular interactions are sufficient to generate, and precisely control, multiphase condensate architectures. By mapping sequence-defined hybridization energies onto macroscopic organization, we provide a predictive design grammar for multiphase condensates. We demonstrate that this interaction energy acts as a master variable and uncover three quantitative design rules: (i) a critical interaction energy threshold sharply delineates complete miscibility from demixing, (ii) above the critical interaction energy, the extent of partitioning scales predictably with interaction energy, and (iii) orthogonal interactions can result in multiple co-existing phases. This scaling behavior, captured by our modified associative polymer theory, provides a quantitative framework linking molecular-level interactions to mesoscale organization. Crucially, each phase retains distinct material properties primarily driven by the dominant component, with >100-fold differences in viscosity across phase boundaries within single droplets. This framework reveals how cells can encode and dynamically regulate multiphase organization by simply tuning interaction affinities (e.g., via post-translational modifications)^73–75^ or component concentrations (e.g., via protein expression)^13^.

A key conceptual insight from our work is that associative phase separation acts as more than a simple concentration mechanism: it fundamentally alters the rules of molecular engagement. By driving macromolecule concentrations above the critical overlap concentration, associative phase separation shifts the system from a dilute solution into a semi-dilute regime where polymer chains interpenetrate, bringing molecules into proximity and enabling new interaction modes that are otherwise rare in dilute solution^76^. In this emergent state, even modest homotypic preferences can overcome the entropy of mixing to drive spatial segregation. This model may have implications for cellular condensates, where enrichment and colocalization may "switch on" new interactions that are inaccessible in the bulk cytoplasm or nucleoplasm^77,78^. Moreover, this principle suggests how compartmentalization might have emerged in early life: before the evolution of lipid membranes, simple polymers with weak, sequence-specific interactions could have created distinct chemical environments through phase separation within condensates, providing primitive cells with a mechanism for spatial organization and functional specialization^79^.

Our demonstration that sequence-specific RNA-RNA interactions can dictate multiphase organization highlights how RNA itself can act as a key architectural component and scaffold complex RNP granules^80,81^. RNA folding, chemical modifications or RNA-binding proteins may allow cells to mask or expose interaction sites and thus modulate partitioning and multiphase architecture. The biophysical insights may also be pertinent to proteins: Many proteins that drive phase separation contain intrinsically disordered regions (IDRs), that can be effectively described as "sticker-and-spacer" polymers, analogous to our patchyDNA^56^. The stickers can be aromatic residues, charged motifs, or other domains that engage in weak, multivalent interactions^14,56^. Our work suggests that subtle variations in the sequence of these IDRs, or post-translational modifications that alter sticker affinity, could be sufficient to drive the segregation of proteins into distinct sub-compartments within a single condensate.

Our work also provides a powerful blueprint for the bottom-up engineering of synthetic soft matter. We can engineer droplets containing coexisting domains with distinct, sequence-encoded material properties. This opens the door to potentially creating synthetic organelles with programmed functions, such as microreactors with sequential reaction chambers or sensors that report on their environment through architectural rearrangements^82,83^. A remarkable feature of these condensates is the existence of sharp, stable interfaces. Rather than passive dividers, these boundaries are quasi-two-dimensional microenvironments with distinct solvation and electrochemical properties^84,85^. Molecules with affinities for both adjoining phases can be enriched at the interface, where reduced dimensionality biases orientation and can enhance catalytic activity. The boundary can also act as a semi-permeable gate, biasing residence times and imposing reaction order as intermediates move between domains.

Our work establishes an equilibrium baseline for multiphase organization, linking molecular interaction energies to mesoscale architecture. In cells, this landscape is elaborated by additional layers of complexity: folding of long RNAs can juxtapose distant interaction sites or occlude them^86^, and active processes, including RNA production and turnover, can drive the system far from equilibrium. Recent work shows that transcriptional flux can set and maintain multiphase order^87^. Integrating these equilibrium principles with energy-consuming processes will be key to creating truly life-like materials and developing a predictive understanding of biological self-organization.

## Author contributions

SM and AJ conceptualized the project and designed the experiments. SM and AN conducted the experiments and analyzed the data. AA and JS developed the theoretical model. AJ and JS secured funding. All authors wrote the manuscript.

## Supporting information

Theory Supplementary Information (Theory SI)

## Acknowledgements

We thank Ofer Kimchi, Marco Todisco, Daniel Stein, Alexander Alfredo Alexander-Katz, and members of Jain lab for helpful discussions. SM and AJ thank Sebastian Coupe and Nikta Fakhri for providing passivated beads, used in micro-rheology experiments. This work was supported by grants from the NIH (AJ: R35GM151111, JS: R01GM141235), Chan Zuckerberg Initiative (AJ: DAF2022-250422), and the David and Lucile Packard Foundation (AJ).

## Notes

### Competing Interest Statement

The authors have declared no competing interest.

