## Supplementary Information (SI) for "Sequence-encoded interactions program internal condensate architecture"

Methods

Supplementary text

Supplementary tables 1-5

Supplementary figures S1-S10

**Methods:**

**Chemicals:**

Nuclease-free water (AM9939) and buffers: Tris pH 7.0 (AM9850G), Tris pH 8.0 (AM9855G) were purchased from Thermo Fisher Scientific, USA. TE buffer was prepared in the laboratory and contained 10 mM Tris, pH 8.0, 0.1 mM EDTA. Fluorescent labeled and unlabeled nucleotides, IDTE pH 7.5 (11-05-01-15) buffer were purchased from IDT, USA. Stock solutions of DNA (250 μM in TE buffer), RNA (250 μM in IDTE pH 7.5), poly-L-lysine (25 mM in nuclease free water) and spermine (100 mM in nuclease free water) were prepared at room temperature. The stock solutions were used fresh or stored at -20°C (DNA, spermine and poly-L-lysine) and -80°C (RNA).

**In-vitro complex coacervation:**

Complex coacervates of a single DNA type were generated by mixing DNA chains and spermine in a PCR tube at room temperature[1]. Complex coacervates of binary DNA mixtures were prepared either by adding spermine to a 1:1 molar mixture of DNA or by mixing pre-prepared coacervate suspensions of the corresponding DNA. Complex coacervates of a single DNA type and binary DNA mixtures were prepared at 10 μM total DNA, 4 mM spermine in 10 mM Tris pH 7.0. Whereas complex coacervates of 3 and 4 DNA mixtures were prepared at 15 μM total DNA, 6 mM spermine in 10 mM Tris pH 7.0 and 20 μM of total DNA, 8 mM spermine in 10 mM Tris pH 7.0, respectively. RNA condensates were prepared by mixing rU60 and r(GGCC) chains with poly-L-lysine (K10) at room temperature at the following buffer condition: 15 μM total RNA, 200 μM K10 in 10 mM Tris pH 7.0, 10 mM $MgCl_2$ and 25 mM NaCl.

**Microscopy**

Freshly prepared coacervate mixtures were transferred to a passivated 384 glass-bottom well plate (Brooks, MatriPlate MGB101-1-2-LG-L) and visualized using a Nikon TE2000E inverted microscope. Samples were illuminated via an Andor Dragonfly 500 spinning disc (Oxford Instruments, USA) system and images were captured using a Andor iXon EM-CCD. 6-FAM, Cy3 and Cy5 labeled samples were imaged using the following combinations of lasers and emission filters: 488 nm with 521 - 38 nm, 561 nm with 600 - 50 nm and 637 nm lasers with 700 – 75 nm, respectively.

**Material properties of coacervates:**

**A. Self-diffusivity of nucleic acids**

Self-diffusivity ($D_{self}$) of DNA chains within multiphase coacervates were measured using fluorescent recovery after photobleaching (FRAP) of Cy3 or Cy5 labeled DNA[2]. A circular spot of diameter d at the center of a compartment was photobleached using a Micro Point pulsed nitrogen-pumped dye laser (405 nm). The FRAP signal ($I$ (t)) was calculated using,

$$I(t) = \frac{B\ (t) - N(t)}{U\ (t) - N(t)} \times \frac{U\ (0) - N(0)}{B\ (0) - N(0)} \qquad (1)$$

Here B (t), U (t) and N (t) denote intensity at bleached spot, unbleached reference spot and background noise, respectively. B (0), U (0) and N (0) are mean (n = 5) intensities before we photobleached the sample. $I$ (t) was normalized ($I_N$ (t)) and the FRAP recovery time ($\tau_{FRAP}$) was calculated by fitting $I_N$ (t) with time (t) using:

$$I_N\ (t) = A\ (1 - e^{\frac{-t}{\tau_{FRAP}}}) \qquad (2)$$

where A is the recovery fraction. Subsequently, $D_{self}$ was calculated using:

$$D_{self} = d^2/4\tau_{FRAP} \qquad (3)$$

### B. Viscosity of coacervates

The viscosity (η) of a phase within coacervates was estimated by measuring mean squared displacement (MSD) of freely diffusing embedded polystyrene beads (radius, $r_{probe}$ = 50 nm, Invitrogen™ FluoSpheres™ F8801)[1,3]. The diffusion coefficient of the beads ($D_{probe}$) was extracted from MSD using the following relation:

$$MSD\ (t) = 4D_{probe}t^{\alpha} + NF \qquad (4)$$

Here, t is the lag time, α is the diffusion exponent and NF is the noise floor at room temperature (T = 22°C). Then, η was calculated using the Stokes Einstein equation:

$$\eta = \frac{k_b T}{6\pi D_{probe} r_{probe}} \qquad (5)$$

**Phase properties of coacervates:**

#### A. Critical salt concentration:

Critical salt concentration (C*) was estimated using two methods[4]. First, coacervate samples (total DNA: 10 μM, spermine = 4 mM in Tris pH 7 = 10 mM) were prepared at 0 – 100 mM NaCl in 5 mM increments (0 mM, 5 mM, 10 mM and so on) and the resulting mixture was imaged. In the first method, we determined C* to be the NaCl concentration at and beyond which no coacervates were observed under the microscope. In the second method, C* was determined from the turbidity of the coacervates. For this, the absorbance of samples at 400 nm was measured using Nanodrop 8000 UV-vis spectrophotometer (Thermofisher, USA). The C* was the NaCl concentration where turbidity (=100-transmittence%) reached a minimum value.

#### B. Soluble DNA fraction:

Soluble DNA fraction is the ratio of DNA concentration in the supernatant phase ($C_{dilute}$) to DNA input concentration (10 μM). To measure $C_{dilute}$, 100 μl coacervate mixtures were prepared in 1.5 ml tubes and centrifuged at 12,000g for 20 minutes. 50 µl of supernatant was collected from each mixture and recentrifuged for 10 min at 12,000g. Then 20 µl of supernatant was collected. $C_{dilute}$ was estimated from the absorbance at 260 nm of the final supernatant phase**.**

#### C. Temperature modulation

To characterize temperature dependent changes in condensates, we reversibly heated and cooled condensate samples in a Cherry Temp temperature control system (Cherry Biotech, France). Typically, we dispensed 10 µl of freshly prepared condensate mixture onto a passivated coverslip (VWR, USA, 48393-251) at room temperature. Then, a temperature controlling chip was placed on top of the sample, and the temperature of the sample was manually controlled using Cherrytemp software. To avoid liquid spillage, the coverslip and the chip were separated by a 500 µm spacer. All accessories and the interfacing software were obtained from the manufacturer.

**Partition coefficient in biphase condensates:**

The partition function (K) of a DNA component was measured using the intensity of fluorescent tagged DNA in the biphase condensates[5]. Samples were prepared by mixing T90 and a patchyDNA (total DNA 10 µM and spermine 4 mM) at desirable compositions. To measure We prepared two sets of samples, each with only one fluorescent probe: one with Cy3 labeled T90 DNA (<1% of T90) and another with only Cy5 labeled patchyDNA (<1% of DNA2), to measure the partition functions of T90 and DNA2, respectively. This minimizes errors originating from bleeding through across fluorescence channels. Partition function was then calculated using the following formula:

$$K = \frac{I_{rich} - I_N}{I_{poor} - I_N} \quad (6)$$

Here, $I_{\text{rich}}$ and $I_{\text{poor}}$ are intensity values averaged over a 4×4-pixel ROI within the phase enriched by the component and in the other phase, respectively. $I_{\text{N}}$ is background noise.

**Supplementary note 1:**

**Design of patchyDNA and patchyRNA**

The patchy nucleic acids were designed to induce a transient network of DNA or RNA chins within the condensate phase[1]. These nucleic acids are single stranded, and all patchyDNA have the same degree of polymerization (90 nucleotides). We used T-90 as backbone to design patchyDNA for two reasons. First, T-90 is canonically inert and cannot form a secondary structure. Second, T-90 spontaneously forms DNA-dense coacervates, where the density of DNA (about 10 mM) is comparable to the overlap concentration (about 1 mM) of T-90. We integrated self-associating sequences or patches to the inert T-90 scaffold to enable inter-DNA interaction within coacervates. The patches are 2 to 8 nucleotide long palindromic sequences (GC, GTAC, GGATCC and so on) that can self-associate at room temperature. We integrated 4 patches per chain, with subsequent patches spaced by long stretches of polyT (15 – 19) to reduce cooperativity[6,7].

The theoretical self-hybridization energy of the patches was predicted using NUPACK (www.nupack.org)[8,9]. For all patches the calculation conditions were the same: 10 mM of patch, 1 M NaCl, NUPACK model DNA04 (allowing dangle and coaxial stacking) at desired temperatures. Here we used patch density similar to DNA density (≈10 mM) within condensates. Since the actual ion environment within DNA/spermine condensates is not known, we used solution conditions that would yield a non-zero value for hybridization for all tested conditions. These conditions may be very different from the actual environment within the coacervate but provide a suitable proxy to compare the self-association energy of patches.

The patchyRNA r(GGCC) was designed by placing four rGGCC patches, on a rU-60 backbone[1]. Successive patches were separated by 12 nucleotide long polyU spacers. Hybridization energies of rGGCC patches were calculated using the following conditions in NUPACK: 40 mM of patch, 1 M NaCl, NUPACK model RNA06 (allowing dangle and coaxial stacking) at desired temperatures.

**Supplementary tables:**

**Supplementary Table 1.**

List of patchyDNA with 4 patches and self-binding interactions of patches at room temperature:

| DNA name | 5' - 3' sequence | patch sequence | ε= -ΔG/RT<br>T = 295 K |
|---|---|---|---|
| T-90 | TTTTTTTTTTTTTTTTTTTTTTTTTTTTTTTTTTTTTTTTTTTTTTTTTTTTTTTTTTT TTTTTTTTTTTTTTTTTTTTTTT TTTTTTTT | - | 0 |
| GC | TTTTTTTTTGCTTTTTTTTTTTTTTTTTTTTTGCTTTTTTTTTTTTTTTTTTTTTGCTT TTTTTTTTTTTTTTTTTTGCTT TTTTTTTT | GC | 5.26 |
| GTAC | TTTTTTTTGTACTTTTTTTTTTTTTTTTT TGTACTTTTTTTTTTTTTTTTTT TGTACTTTTTTTTTTTTTTTTTT TGTACTTTTTTTTT | GTAC | 8.4 |
| GATC | TTTTTTTTGATCTTTTTTTTTTTTTTTTT TGATCTTTTTTTTTTTTTTTTTT TGATCTTTTTTTTTTTTTTTTTT TGATCTTTTTTTTT | GATC | 8.4 |
| AATT | TTTTTTTTAATTTTTTTTTTTTTTTTTTTTT TAATTTTTTTTTTTTTTTTTTTTT TAATTTTTTTTTTTTTTTTTTTTT TAATTTTTTTTTTT | AATT | 6.84 |
| GGCC | TTTTTTTTGGCCTTTTTTTTTTTTTTTTT TGGCCTTTTTTTTTTTTTTTTTT TGGCCTTTTTTTTTTTTTTTTTT TGGCCTTTTTTTTT | GGCC | 12.54 |
| CGCG | TTTTTTTTCGCGTTTTTTTTTTTTTTTTT TCGCGTTTTTTTTTTTTTTTTTT TCGCGTTTTTTTTTTTTTTTTTT TCGCGTTTTTTTTT | CGCG | 14.04 |
| GCGC | TTTTTTTTGCGCTTTTTTTTTTTTTTTTT TGCGCTTTTTTTTTTTTTTTTTT TGCGCTTTTTTTTTTTTTTTTTT TGCGCTTTTTTTTT | GCGC | 14.09 |
| GAATTC | TTTTTTTGAATTCTTTTTTTTTTTTTTTTT GAATTCTTTTTTTTTTTTTTTTTT GAATTCTTTTTTTTTTTTTTTTTT GAATTCTTTTTTTT | GAATTC | 12.95 |
| GTTAAC | TTTTTTTGTTAACTTTTTTTTTTTTTTTTT GTTAACTTTTTTTTTTTTTTTTTT GTTAACTTTTTTTTTTTTTTTTTT GTTAACTTTTTTTT | GTTAAC | 12.95 |
| CCTAGG | TTTTTTTCCTAGGTTTTTTTTTTTTTTTTT CCTAGGTTTTTTTTTTTTTTTTTT CCTAGGTTTTTTTTTTTTTTTTTT CCTAGGTTTTTTTT | CCTAGG | 15.08 |
| GGATCC | TTTTTTTGGATCCTTTTTTTTTTTTTTTTT GGATCCTTTTTTTTTTTTTTTTTT GGATCCTTTTTTTTTTTTTTTTTT GGATCCTTTTTTTT | GGATCC | 15.69 |
| CCATGG | TTTTTTTCCATGGTTTTTTTTTTTTTTTTT CCATGGTTTTTTTTTTTTTTTTTT CCATGGTTTTTTTTTTTTTTTTTT CCATGGTTTTTTTT | CCATGG | 16.22 |
| CGATCG | TTTTTTTCGATCGTTTTTTTTTTTTTTTTT CGATCGTTTTTTTTTTTTTTTTTT CGATCGTTTTTTTTTTTTTTTTTT CGATCGTTTTTTTT | CGATCG | 17.19 |
| GGGCCC | TTTTTTTGGGCCCTTTTTTTTTTTTTTTTT GGGCCCTTTTTTTTTTTTTTTTTT GGGCCCTTTTTTTTTTTTTTTTTT GGGCCCTTTTTTTT | GGGCCC | 19.83 |
| GGAATTCC | TTTTTTGGAATTCCTTTTTTTTTTTTTTTTGGAATTCCTTTTTTTTTTTTTTTT GGAATTCCTTTTTTTTTTTTTTTT GGAATTCCTTTTTTT | GGAATTCC | 20.24 |
| GCGCGC | TTTTTTTGCGCGCTTTTTTTTTTTTTTTTT GCGCGCTTTTTTTTTTTTTTTTTT GCGCGCTTTTTTTTTTTTTTTTTT GCGCGCTTTTTTTT | GCGCGC | 22.91 |

**Supplementary Table 2.**

Self-binding interaction of DNA patches at different temperatures:

| | $\varepsilon = -\Delta G/RT$ | | | | |
|---|---|---|---|---|---|
| temperature (K) | GAATTC | GTTAAC | GGCC | GGATCC | GGAATTCC |
| 293 | 13.40 | 13.41 | 12.83 | 16.14 | 20.87 |
| 298 | 12.28 | 12.26 | 12.11 | 15.01 | 19.29 |
| 303 | 11.18 | 11.16 | 11.39 | 13.91 | 17.75 |
| 308 | 10.13 | 10.10 | 10.7 | 12.86 | 16.27 |
| 313 | 9.10 | 10.68 | 10.03 | 11.82 | 14.82 |
| 318 | 8.11 | 8.06 | 9.37 | 10.82 | 13.42 |

### Supplementary Table 3.

Cross interaction between patches at room temperature

| | ΔG (Kcal/mol) | | | |
|---|---|---|---|---|
| patch 1 | homotypic interaction | patch 2 | homotypic interaction | heterotpic interaction |
| GGATCC | -9.21 | GGGCCC | -11.64 | -4.8 |
| GGATCC | -9.21 | GGAATTCC | -11.88 | -4.96 |
| GGGCCC | -11.64 | GGAATTCC | -11.88 | -4.8 |

### Supplementary Table 4.

List of RNA

| RNA name | 5' - 3' sequence | patch sequence | $\varepsilon_{RNA}$=-ΔG/RT at T = 295 K |
|---|---|---|---|
| rU-60 | UUUUUUUUUUUUUUUUUUUUUUUUUUUUUUUUUUUUUUUUUUUUUUUUUUUUUUUUUUUU | - | 0 |
| r(GGCC) | UUUUGGCCUUUUUUUUUUUUGGCCUUUUUUUUUUUUGGCCUUUUUUUUUUUUGGCCUUUU | GGCC | 16.73 |

### Supplementary Table 5.

Self-binding interaction of RNA patches at different temperatures:

| | εRNA= -ΔG/RT |
|---|---|
| temperature (K) | rGGCC |
| 298 | 16.01 |
| 303 | 14.95 |
| 308 | 14.06 |
| 313 | 13.03 |
| 318 | 12.03 |

**Supplementary figures:**

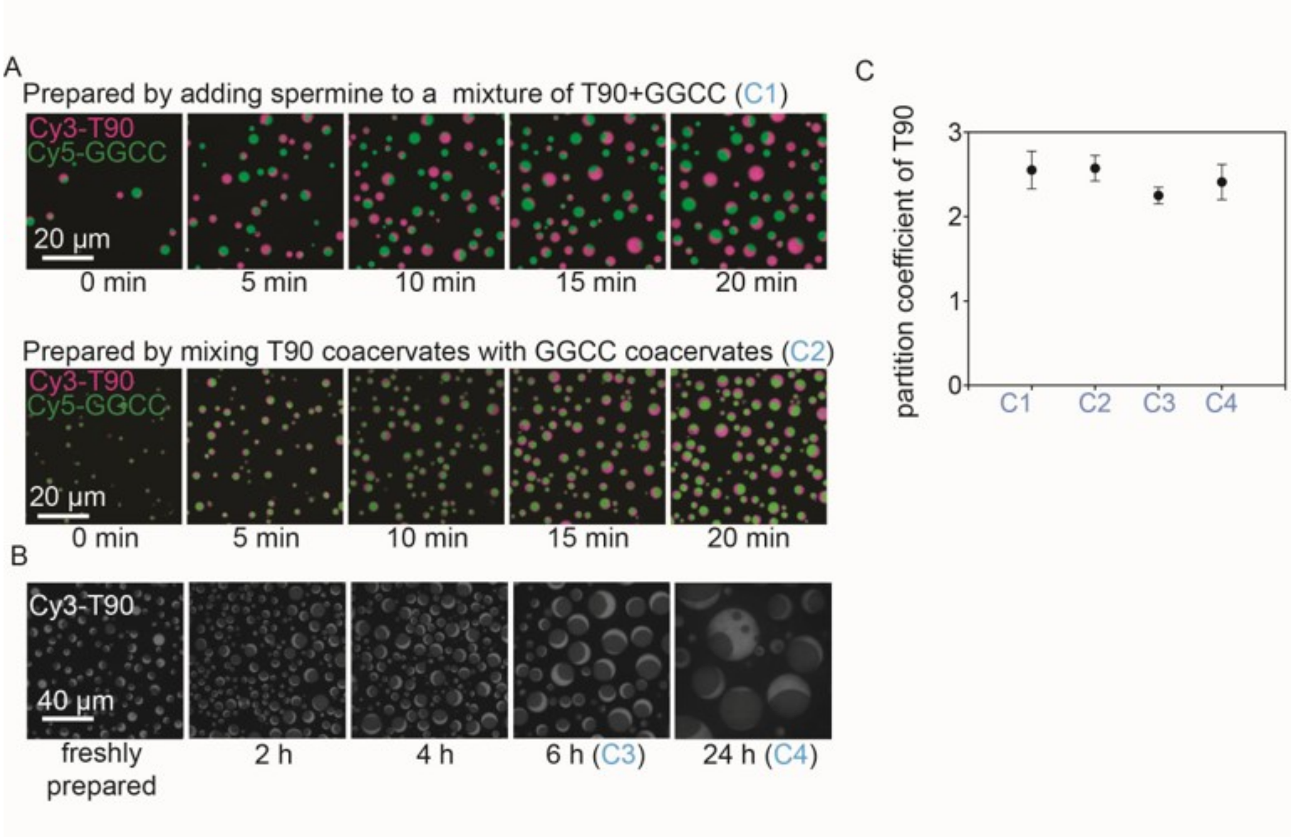


**Figure S1: Two-phase condensates are preparation independent and remain stable with time:** Representative fluorescent micrographs of T90+GGCC/spermine condensates prepared by indicated methods (A) and at different times post preparation (B). (C) Plot showing that partition coefficient of T90 did not perturb with preparation method or time. All condensates were prepared at the following condition: T90=5 µM, GGCC=5µM, spermine= 4 mM in 10 mM Tris pH 7.0 at 22 $^{o}$C.

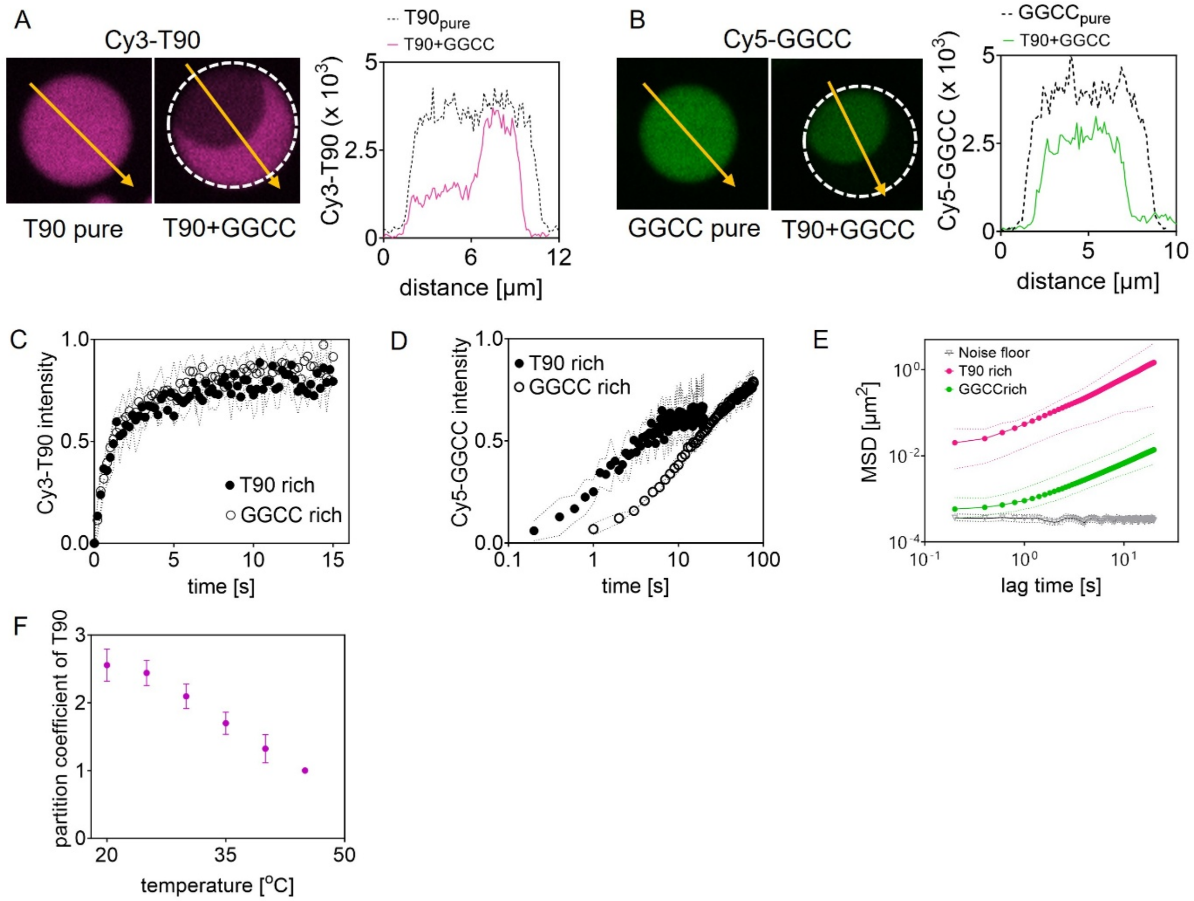


**Figure S2: Characterization of T90+GGCC/spermine two-phase condensates:** (A-left) Representatives fluorescent micrographs and (A-right) corresponding intensity traces, comparing T90 density in two-phase condensates with pure T90 condensate. (B) Same as A but for GGCC. FRAP recovery curves of (C) Cy3-T90 and (D) Cy5-GGCC in the indicated compartments in T90+GGCC/spermine two-phase condensates. (E) Plot comparing MSD traces of embedded beads (radius = 50nm) in the indicated compartments with the noise floor at room temperature. (F) Partition coefficients of T90 in T90+GGCC/spermine condensates as a function of temperature. Here all condensates were prepared at the following condition: DNA=10μM, spermine= 4 mM in 10 mM Tris pH 7.0 at 22 $^{o}$C. All two-phase condensates were prepared at 1:1 molar ratio of T90 and GGCC.

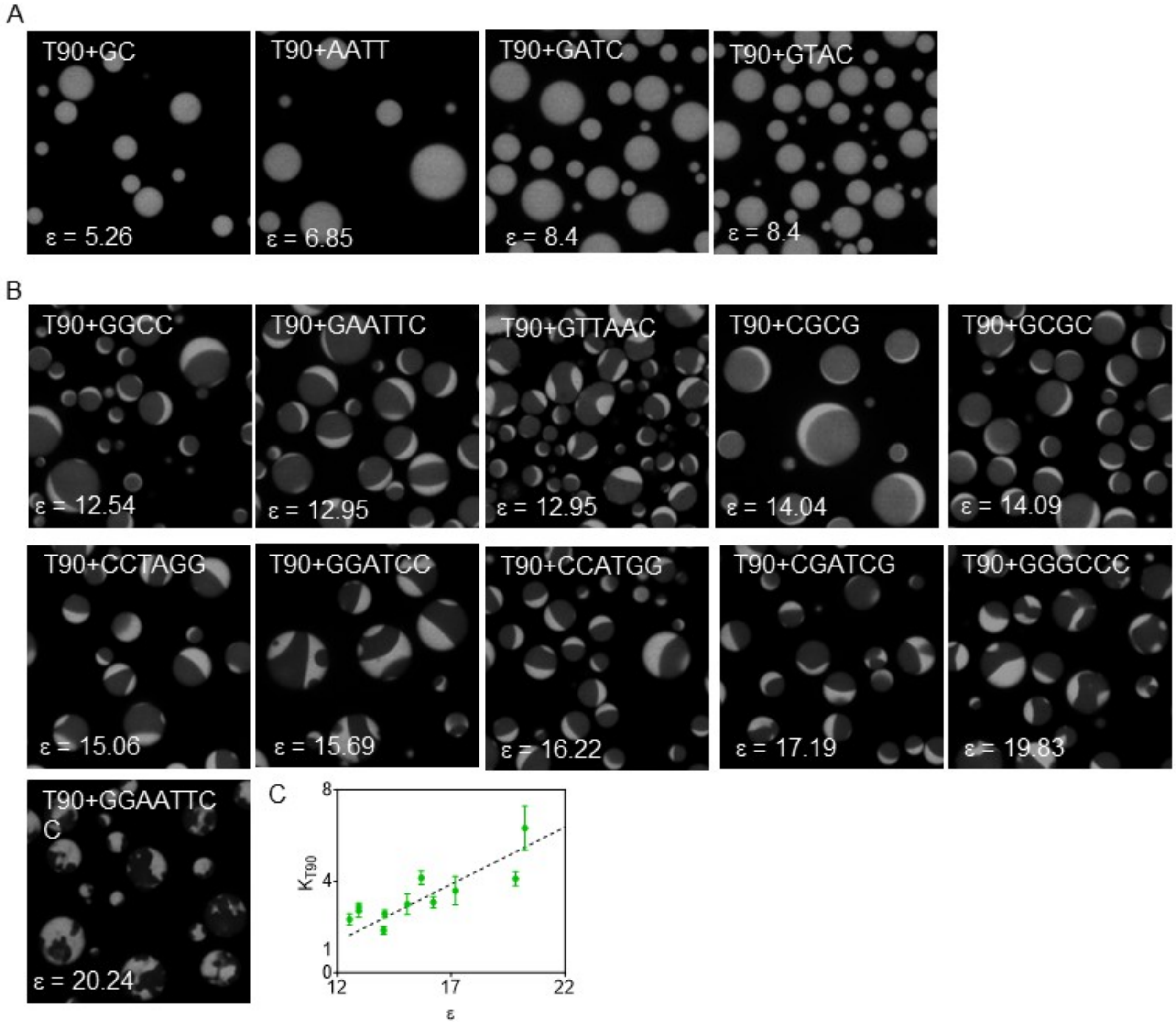


**Figure S3**: **Single to two-phase DNA/spermine condensates:** (A-B) Representative micrographs showing condensates of T90+patchyDNA/spermine with indicated patches. ε represents hybridization strength of patches at room temperature (22 ºC). (C) Plot showing partition coefficient of T90 ($K_{T90}$) in T90+patchyDNA/spermine condensates formed at 22 ºC. The black dotted line is for visual purpose only and has a slope of 0.5. All condensates were prepared at 1:1 molar ratio of T90 and patchyDNA at the following conditions: total DNA = 10µM, spermine = 4 mM in 10 mM Tris pH 7.0 at 22 ºC. All images have a size of 30 × 30 µm$^2$.

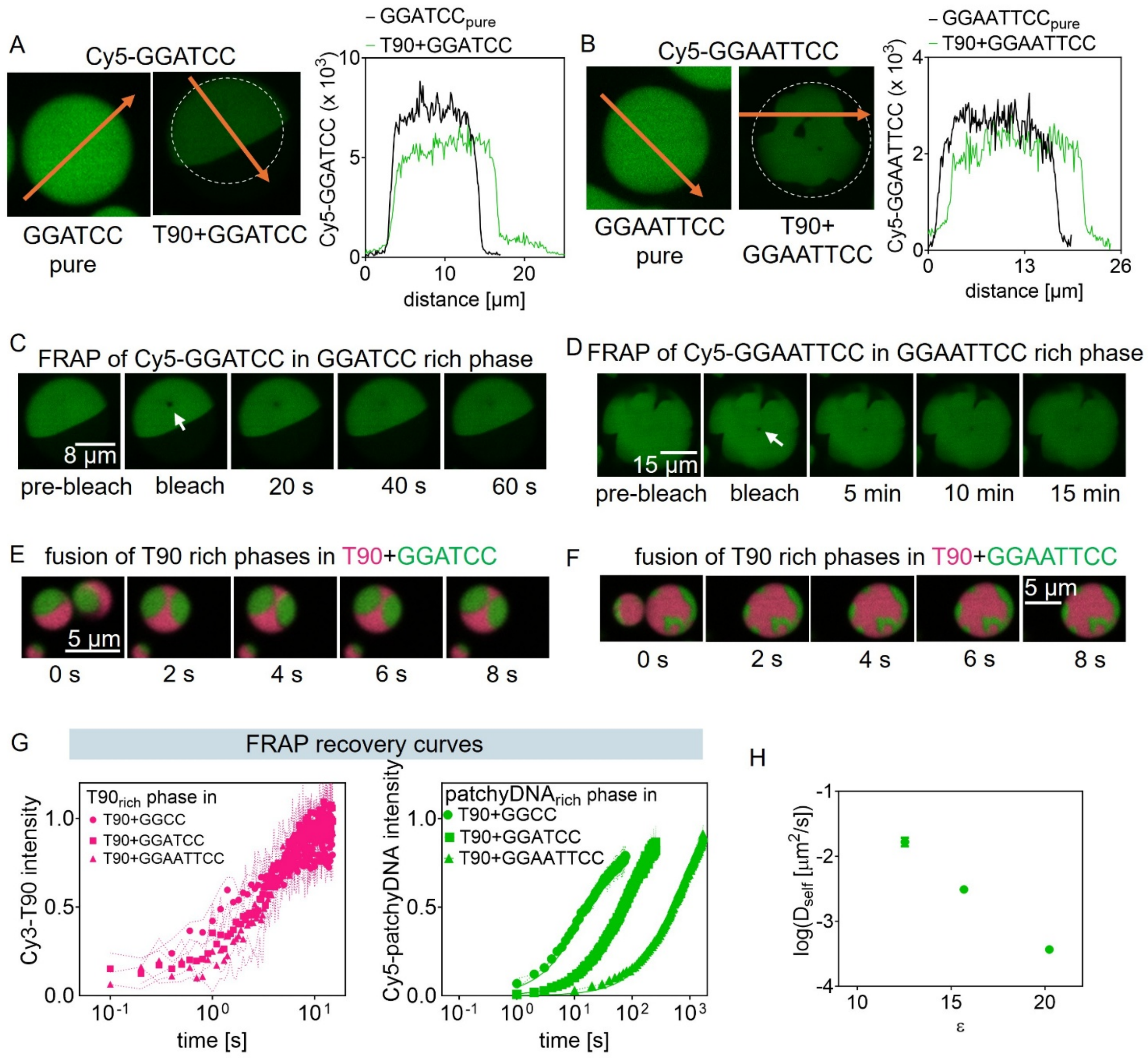


**Figure S4: Characterization of T90 rich and patchyDNA rich phases across various T90+patchyDNA/spermine condensates:** (A-left) Representatives fluorescent micrographs and (A-right) corresponding intensity traces, comparing GGATCC density in T90+GGATCC/spermine condensates with pure GGATCC condensate. (B) Same as A but for GGAATTCC. (C) Representative fluorescent micrographs showing FRAP recovery of GGATCC in GGATCC rich phase in a T90+GGATCC/spermine condensate. (D) Same as C, but for GGAATTCC. Representative fluorescent micrographs showing fusion of T90 rich phases in (E)

T90+GGATCC/spermine condensates and (F) T90+GGAATTCC/spermine condensates. (G) FRAP recovery curves of T90 and patchyDNA chains in the indicated compartments. (H) Plot showing ε dependent self-diffusivity ($D_{self}$) of patchyDNA chains in their corresponding rich compartments within T90+patchyDNA/spermine condensates. Here all condensates were prepared at the following condition: DNA=10μM, spermine= 4 mM in 10 mM Tris pH 7.0 at 22 $^{o}$C. All two-phase condensates were prepared at 1:1 molar ratio of T90 and GGCC.

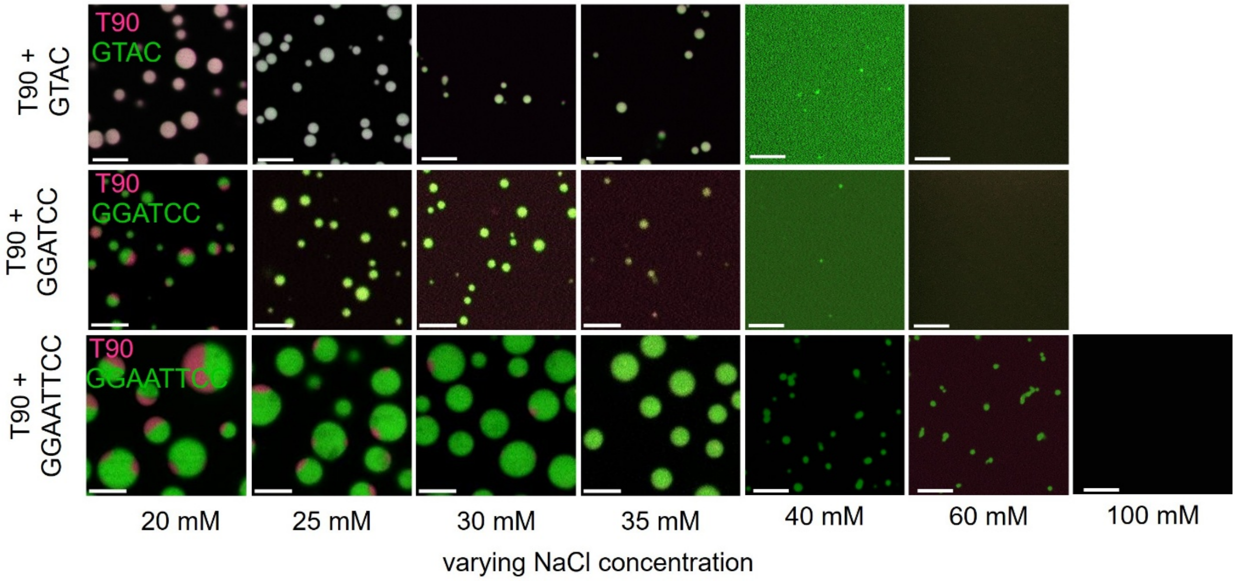


**Figure S5: Phase diagram with NaCl:** Representative fluorescent micrographs showing NaCl mediated dissolution of T90+patchyDNA/spermine condensates. All condensates were prepared at 1:1 molar ratio of T90 and patchyDNA at the following conditions: total DNA = 10µM, spermine = 4 mM in 10 mM Tris pH 7.0 at 22 °C. The scale bars have a value of = 5 µm.

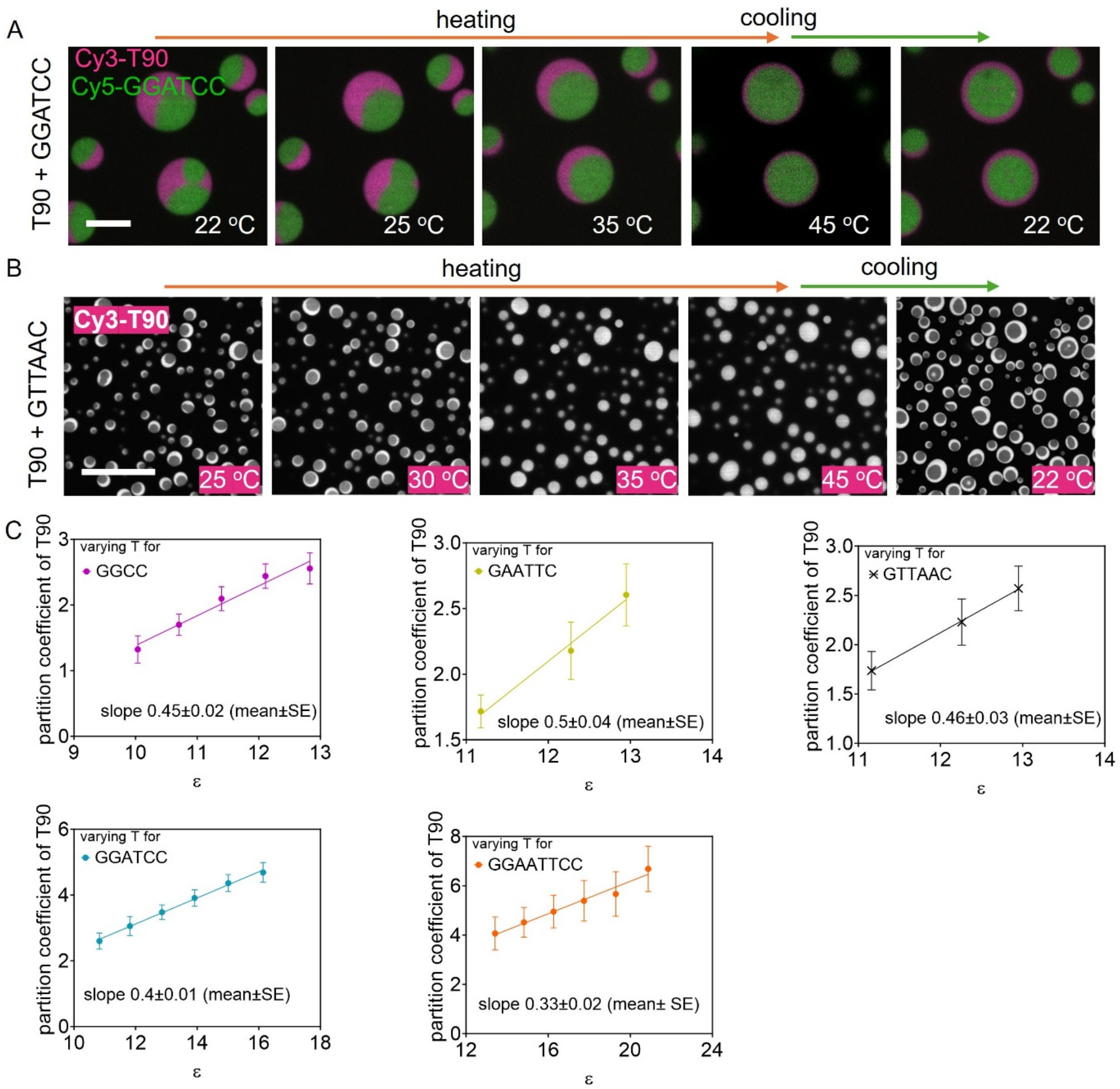


**Figure S6: Temperature dependent phase-separation:** Representative fluorescent micrographs showing that, two-phase condensates of T90+GTTAAC/spermine (A) reversibly dissolve and segregate with temperature, whereas T90+GGATCC/spermine (B) condensates maintain bi-phase morphology even at 45 $^{o}$C. (C) partition coefficients of T90 in T90+patchyDNA/spermine condensates of the indicated patchyDNA. Here ε was varied by temperature. The straight lines in C represent linear fits to the data with the indicated slopes. Scale bars in A and B have values 5 μm and 30 μm, respectively.

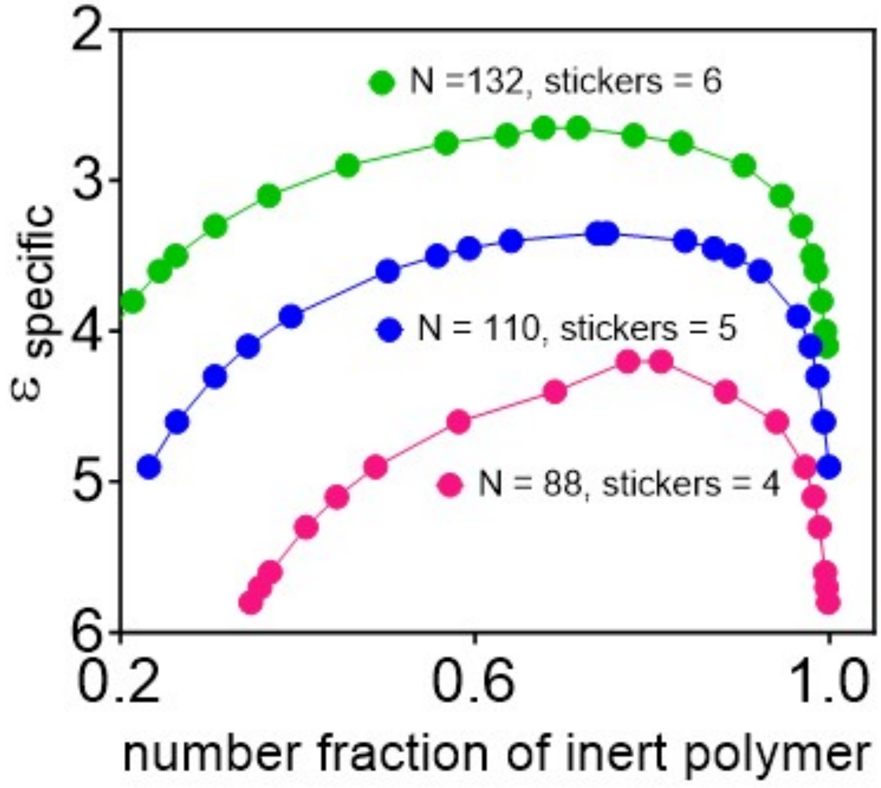


**Figure S7: Theory phase diagram as a function of valence:** Phase diagram for demixing of phases in a binary mixture of an inert polymer and a sticky polymer with indicated chain lengths (N).

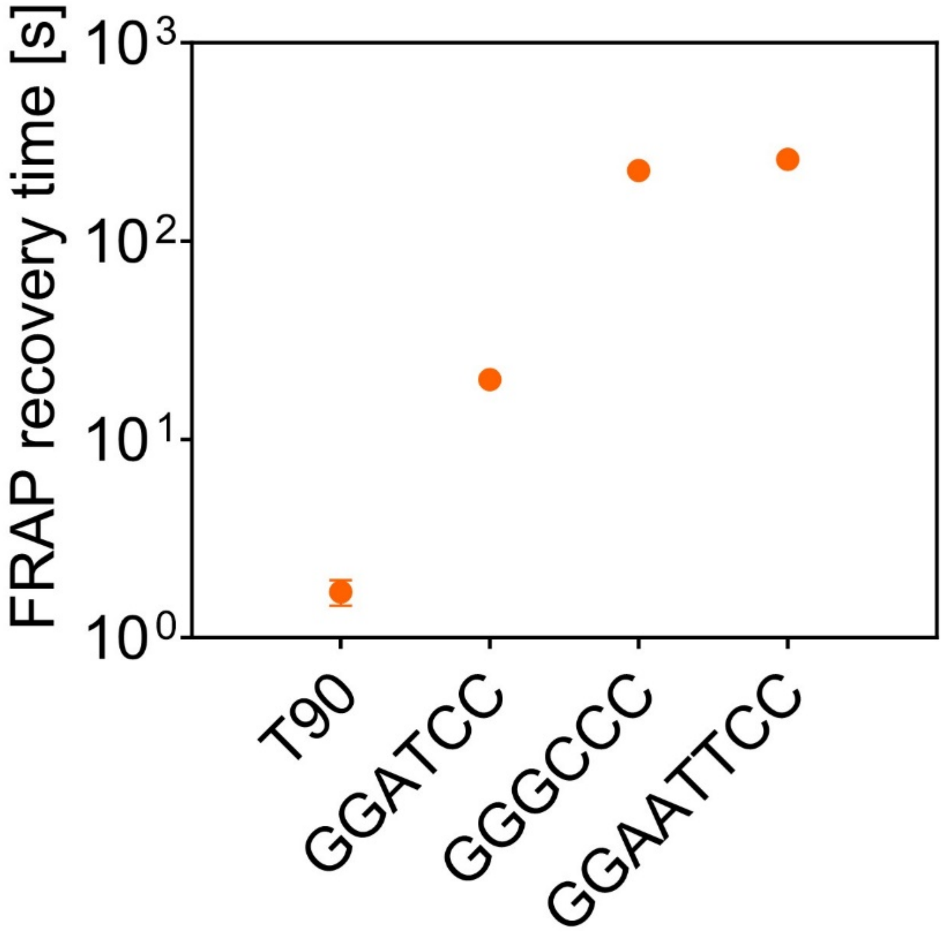


**Figure S8: Co-exiting phases within multiphase condensates exhibit sequence dependent material properties:** FRAP recovery time of indicated DNA in their respective rich phases within four phase condensates of T90+GGATCC+GGAATTCC+GGGCCC/spermine.

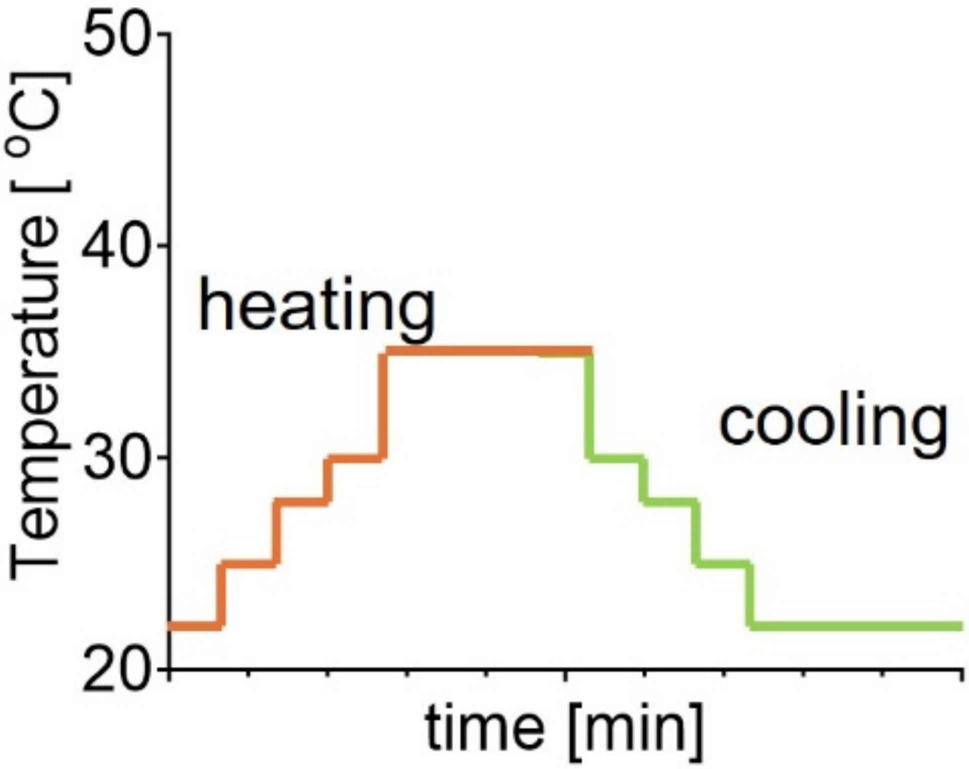


**Figure S9: Heat treatment of RNA condensates:** Plot showing step-temperature protocol used for heating and cooling rU60+r(GGCC)/poly-L-lysine condensates.

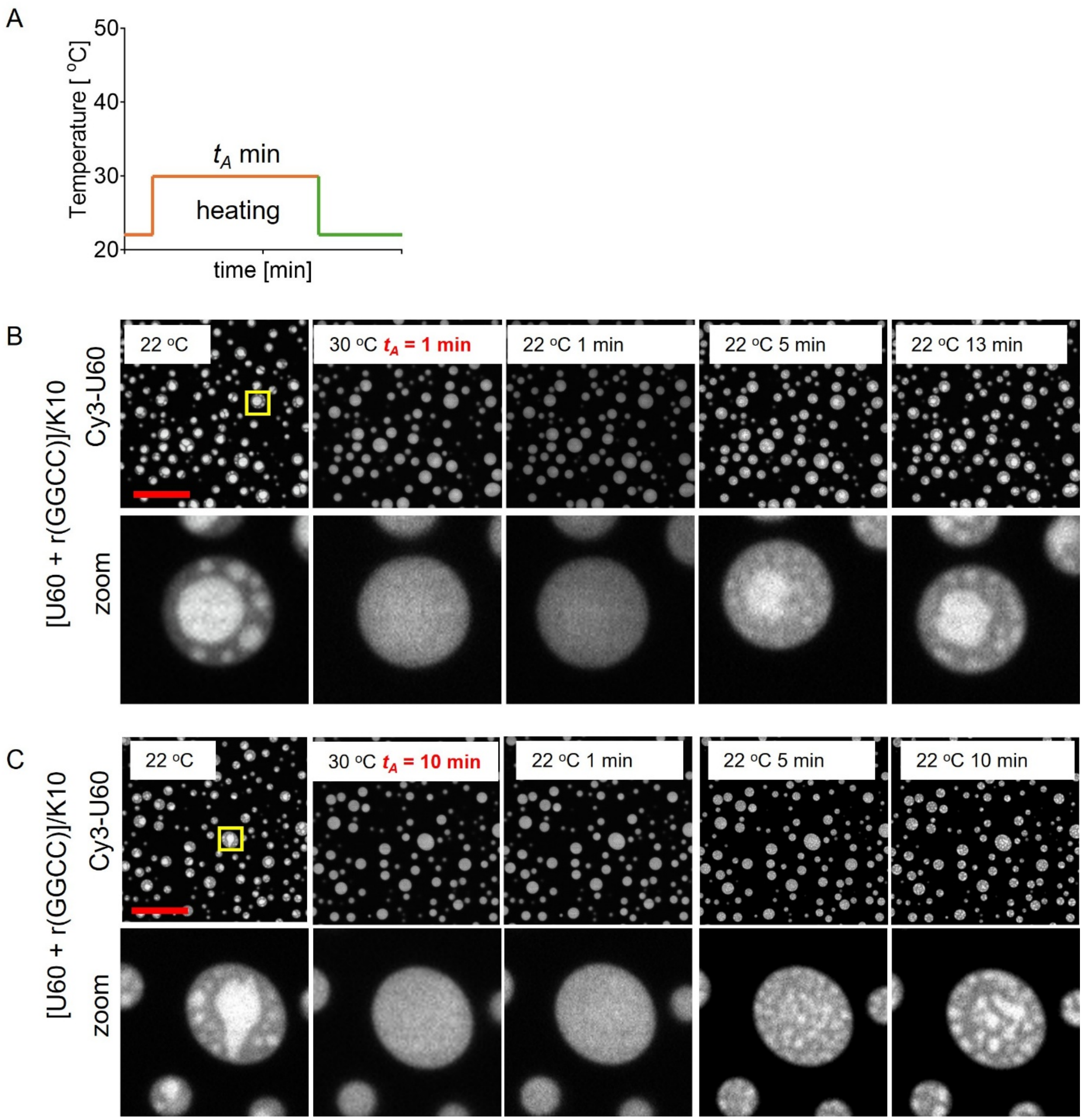


**Figure S10: Melt memory in RNA condensates:** (A) Plot showing step-temperature protocol used to test melt memory in two-phase condensates of rU60+r(GGCC)/poly-L-lysine (K10). Typically, two-phase RNA samples were heated from 22°C to 30°C. Then the samples were annealed for different times ($t_A$) at 30 °C before quenching back to 22°C. (B-C) Representative fluorescent micrographs showing morphology of rU60+r(GGCC)/K10 condensates at indicated temperatures. For a short annealing time ($t_A$) = 1 min, upon quenching to room temperature, rU60 rich phases nucleated predominantly at the sites where rU60 rich compartments were present before heating. However, for a long annealing time ($t_A$) = 10 min, the condensates lost the sample memory and upon cooling rU60 rich phases nucleated randomly throughout the condensates. The scale bars in B-C have a value of 15 μm.
