## Supplementary material for "Sequence-encoded interactions program internal condensate architecture": Theory Supplementary Information (Theory SI)

### Calculation of phase diagram

Let there be a system composed of two different types of polymers. In our case, we are specifically focusing on polymers which can be modeled as sticker-spacer chains (if there are no stickers, the chain is then just a long spacer). Stickers are composed of beads which participate in the formation of homotypic or heterotypic bonds. Spacers are inert beads between two subsequent stickers. Two different types of polymer chain will have different sticker-spacer profile. Our goal is to determine if a condensate containing two chain types (hitherto identified as A-type and B-type) will remain mixed or separate into two distinct phases. Formally this can be done by taking the convex hull of the three component free energy (A, B, and solvent) [1]. Here we make the simplifying assumption that the separation of the dense phase into A-rich and B-rich regions will minimally perturb the composition of the dilute phase. In this pseudo-two-phase approximation the free energy of the two dense phases is

$$F_{tot} = n_1 f_{chain}(\psi_1) + n_2 f_{chain}(\psi_2) - \lambda_1(n_1 + n_2 - n_{tot}) - \lambda_2(\psi_1 n_1 + \psi_2 n_2 - n_s) \quad (1)$$

where $n_1$ and $n_2$ refer to the total number of chains in each phase, $n_{tot} = n_1 + n_2$ is the number of chains, and $f_{chain}$ is the free energy per chain. The free energy depends on the composition of a given phase, which is captured by the variable $\psi = \dfrac{C_A}{C_A + C_B} = \dfrac{n_A}{n_A + n_B}$, where $c_i(n_i)$ denote the concentration(number) of $A$ or $B$ molecules and $n_s = \psi_1 n_1 + \psi_2 n_2$ is the total number of $A$ chains. It's also important to note that $n_1 + n_2 = n_A + n_B$. The Lagrange multipliers $\lambda_1$ and $\lambda_2$ thus constrain the total number of molecules and average composition of the dense phase. Note that, unlike the usual double tangent derivation, we do not apply a volume constraint because the dense phases can expand or contract within the engulfing dilute phase.

Minimizing with respect to $\psi_1$, $\psi_2$, $n_1$ and $n_2$, we arrive at equations analogous to the usual double tangent construction

$$\frac{\partial F_{tot}}{\partial \psi_1} = n_1 f'_{chain}(\psi_1) - \lambda_2 n_1 = 0 \tag{2}$$
$$\frac{\partial F_{tot}}{\partial \psi_2} = n_2 f'_{chain}(\psi_2) - \lambda_2 n_2 = 0 \tag{3}$$
$$\frac{\partial F_{tot}}{\partial n_1} = f_{chain}(\psi_1) + \lambda_1 - \lambda_2 \psi_1 = f_{chain}(\psi_1) + \lambda_1 - \psi_1 f'_{chain}(\psi_1) = 0 \tag{4}$$
$$\frac{\partial F_{tot}}{\partial n_2} = f_{chain}(\psi_2) + \lambda_1 - \lambda_2 \psi_2 = f_{chain}(\psi_2) + \lambda_1 - \psi_2 f'_{chain}(\psi_2) = 0 \tag{5}$$

where $\lambda_1$ is the y-intercept and $\lambda_2$ is the slope.

Our phase free energies are based on the Semenov-Rubenstein model [2]. This is a mean-field theory describing attractive "sticker" beads mixed with non-attractive (or weakly attractive) "spacer" beads [2] [3]. In the Semenov and Rubenstein theory, the driving force for condensation comes from the terms describing the free energy of sticker-sticker bonds along with the combinatorial entropy of choosing the sticker pairs. This attraction is balanced against the mixing entropy cost of concentrating the molecules and the repulsion between the spacer beads. Here we use Semenov-Rubinstein expressions for the bonding and mixing contributions, but use an alternate form for the spacer-spacer interactions to account for the complexities of the spermine-mediated spacer-spacer interactions.

The sticker-sticker contribution to the free energy is given by

$$F_{\text{bind}} = P\epsilon - \ln \Omega_P \tag{6}$$

where $\epsilon$ is the free energy of a sticker-sticker bond, $P$ is the number of bonds formed, and $\Omega$ is the number of ways to arrange $P$ bonds between $M$ stickers in the system. The combinatorial factor is given by

$$\Omega_P = \left(\frac{v_0}{V}\right)^P \frac{M!}{(M-2P)!\,P!\,2^P} \tag{7}$$

This expression can be understood as follows. There are $M!/[(M-2P)!(2P)!]$ ways to choose $2P$ stickers that are involved in bonds. To form the first bond there are $2P$ options for the first sticker. While there are $2P-1$ options for the second sticker, a bond can only form if the second sticker is sufficiently close to the first. If the volume accessible to a sticker is $v_0$, the number of stickers available to the first sticker within that volume is $(v_0/V)(2P-1)$, where $V$ is the volume of the system. This gives a total of $2P(v_0/V)(2P-1)$ options for the first bond. Similarly, the number of choices for the second bond is $(2P-2)(v_0/V)(2P-3)$. Repeating this argument for all $P$ bonds gives a combinatorial factor of $(2P)!(v_0/V)^P$. This counting needs to be corrected by the $P!$ ways that the same bonds can be selected in a different order and the $2^P$ ways that the bond pairs can be reversed. Putting these contributions together gives Eq.7. Eq.7 increases for more condensed states because small values of $V$ increase the number of bonding options accessible to the stickers. Note that neither term in Eq.6 accounts for the constraint that certain stickers are constrained to be nearby due to the spacer connecting them. However, neglect of this constraint gives the correct answer provided that the spacers are long enough to disfavor consecutive, zipper-like bonds between the same two molecules [4] and the spacers are well mixed (which is ensured when the spacers are uniform length)[5].

Applying Stirling's approximation $\ln(n!) \simeq n \ln n - n$, to Eq.6 we find,

$$\begin{aligned} F_{bind} &= P\epsilon - \ln \Omega_P \\ &= P\epsilon - \{P \ln v_0 - P \ln 2V - (M-2P)\ln(M-2P) - P\ln P + M \ln M - P\} \qquad (8) \\ &= P\left(\epsilon - \ln \frac{v_0}{2PV}\right) + (M-2P)\ln(M-2P) + P - M\ln M \qquad (9) \end{aligned}$$

This becomes with $m = \frac{M}{V}$ and $p = \frac{P}{V}$,

$$f_{bind} = F_{bind}/V \tag{10}$$
$$= p\left(\epsilon - \ln\frac{v_0}{2p}\right) + (m-2p)\ln(m-2p) + p - m\ln m$$

Minimizing $f_{bind}$ with respect to p, we get,

$$0 = \epsilon - \ln\frac{v_0(m-2p)^2}{2p} \tag{11}$$

$$e^{\epsilon} = \frac{v_0(m-2p)^2}{2p} \tag{12}$$

The solution for optimized $p$ is,

$$\frac{p}{m} = \frac{P}{M} = \frac{k+2}{4} - \frac{\sqrt{(k+2)^2-4}}{4} \tag{13}$$

Where, $k = e^{\epsilon - \ln(mv_0)}$.

Our treatment of the spacer-spacer interaction differs from the Semenov/Rubinstein theory, in which the spacers were captured by two-body and three-body virial terms. In our DNA condensates the spacers repel by electrostatic and excluded volume interactions while attracting via spermine-mediated salt bridges. The balance between these contributions gives an optimal concentration $C_0$ for stickerless DNA. Deviations from this value, driven by sticker-sticker interactions, incur a free energy penalty that is quadratic to lowest order

$$f_{repulsion} = \alpha(C - C_0)^2 \tag{14}$$

Within the mean-field treatment the sticker term depends only on the concentration of stickers $m$, while the spacer term depends only on the polymer concentration. This separation between the sticker and spacer terms allows us to treat a mixture of chains that we refer to as type "A" and type "B" by adding the sticker and spacer contributions of each type to the local concentration. We define $n_A$ as the number of $A$ chains and $n_B$ as the number of $B$ chains with corresponding concentrations $C_A = n_A/V$ and $C_B = n_B/V$. The sticker concentration is $m = v_A C_A + v_B C_B$, where $v_A$ and $v_B$ are the number of stickers (valence) per chain. Since we are interested in chains of uniform length, Eq. 14 can account for mixtures with the substitution $C = C_A + C_B$. Finally, we need to separately account for the mixing entropy of the two types by writing

$$F_{mix} = -\ln \Omega_{mix} = n_A \ln \frac{n_A}{V} + n_B \ln \frac{n_B}{V} - (n_A + n_B) + Const. \tag{15}$$

$$f_{mix} = F_{mix}/V = C_A \ \ln C_A + C_B \ \ln C_B - (C_A + C_B) \tag{16}$$

We ignore any constant as they are not relevant in our calculation of phase composition. Collecting all the terms, we have

$$f = (F_{bind} + F_{repulsion} + F_{mix})/V \tag{17}$$

$$= p\left(f - \ln \frac{v_0}{2p}\right) + (m - 2p)\ln(m - 2p) + p - m \ln m$$

$$+ \alpha\,(C_A + C_B - C_0)^2 + C_A \ \ln C_A + C_B \ \ln C_B - (C_A + C_B) \tag{18}$$

In order to apply Eqs. 2-5 we need the free energy per molecule. This is given by $f_{chain} = f/(C_A + C_B)$.

$$f_{chain} = \frac{F_{bind}}{n_A + n_B} + \alpha \frac{C_0^2}{C}\left(\frac{C}{C_0} - 1\right)^2 + (\psi_A \ \ln \psi_A + \psi_B \ \ln \psi_B + \ln C) \tag{19}$$

$$f_{chain} = \frac{F_{bind}}{n_A + n_B} + \alpha \frac{C_0^2}{C}\left(\frac{C}{C_0} - 1\right)^2 + \psi \ \ln \psi + (1 - \psi) \ \ln (1 - \psi) + \ln C \tag{20}$$

Due to Eqs. 2-5, the system will phase separate whenever Eq. 20 has a region with a negative second derivative. In these cases, the compositions of the two phases, $\psi_1$ and $\psi_2$, can be determined from the common tangent construction.

#### Phase separation in binary mixtures of an inert and a sticky polymer

To understand the biophysical principle governing phase separation in mixtures of T90 and patchyDNA condensates, we calculated theoretical phase diagram for de-mixing in a binary polymer blend. We used an inert polymer and a sticky polymer to generate the theoretical phase diagram as they closely match the molecular architecture of T90 and patchyDNA, respectively. Here, both polymers have the same length of 88 beads. The inert polymer is a linear chain of spacer beads only. Whereas, the sticky polymer has 4 stickers per chain and successive stickers are spaced by 19 spacer beads. Each sticker is 3 beads long and stickers from the same or different chains can form bonds via specific interaction ($\epsilon_{specific} = \epsilon/-kT$, k is Boltzmann's constant and T is temperature). In our calculations, we varied $\epsilon_{specific}$ between 3 and 6. Notably, a spacer bead can interact with other inert beads or stickers via non-specific interactions, however the magnitude of such interactions are negligible compared to $\epsilon_{specific}$.

#### Computational Scheme

To find how immiscibility changes with binding affinity between stickers we follow the following computational steps:

- Write the free energy as a function of $\psi$ and $\phi$. We do this by writing the concentration in terms of volume fraction $\phi = N_b b^3/V$, where $N_b$ is the total number of beads of the system and $b$ is a microscopic length scale. Since we treat $b$ as a constant parameter, it's actual value is not important for our overall scheme. $C$ then can then be written in terms of $C = \frac{n}{V} = \frac{n\phi}{N_b b^3}$.

- For a given $\psi$, we then find the optimized $\phi$ for which the free energy is minimum. We create a table of $\{\psi, \phi\}$ for $\psi-$values between 0 and 1. We then evaluate the free energy based on this table.

- We then construct the double tangent from the free energy vs $\psi$ diagram. We use the data from double tangent $\psi_1$ and $\psi_2$ to construct the final phase diagram.

**Partition coefficient calculation**

We calculated partition coefficient of the inert polymer ($K_{inert}$) to characterize miscibility in binary mixtures of an inert and a sticky polymer using the following formula,

$$K_{inert} = \frac{\psi_{rich}}{\psi_{poor}} \tag{21}$$

where $\psi_{rich}$ and $\psi_{poor}$ are number fraction of the inert polymer in the inert polymer rich and sticky polymer rich phase, respectively. When two chains are homogeneously mixed, $K_{inert} = 1$, and for de-mixed phases $K_{inert} > 1$.
